# The Effects of Quantitative Trait Architecture on Detection Power in Artificial Selection Experiments

**DOI:** 10.1101/672683

**Authors:** R. Nicolas Lou, Nina O. Therkildsen, Philipp W. Messer

**Author notes:** Corresponding authors: RNL, PWM.

## Abstract

Evolve and resequence (E&R) experiments, in which artificial selection is imposed on organisms in a controlled environment, are becoming an increasingly accessible tool for studying the genetic basis of adaptation. Previous work has assessed how different experimental design parameters affect the power to detect the quantitative trait loci (QTLs) that underlie adaptive responses in such experiments, but so far there has been little exploration of how this power varies with the genetic architecture of the evolving traits. In this study, we use forward simulation to build a realistic model of an E&R experiment in which a quantitative polygenic trait experiences a short, but strong, episode of truncation selection. We study the expected power for QTL detection in such an experiment and how this power is influenced by different aspects of trait architecture, including the number of QTLs affecting the trait, their starting frequencies, effect sizes, clustering along a chromosome, dominance, and epistasis patterns. We show that all of the above parameters can affect allele frequency dynamics at the QTLs and linked loci in complex and often unintuitive ways, and thus influence our power to detect them. One consequence of this is that existing detection methods based on models of independent selective sweeps at individual QTLs often have lower detection power than a simple measurement of allele frequency differences before and after selection. Our findings highlight the importance of taking trait architecture into account in designing studies of molecular adaptation with temporal data. We provide a customizable modeling framework that will enable researchers to easily simulate E&R experiments with different trait architectures and parameters tuned to their specific study system, allowing for assessment of expected detection power and optimization of experimental design.

## INTRODUCTION

Artificial selection experiments can provide insights into the mechanisms that allow populations to adapt to strong selection pressures (Hill and Caballero 1992; Fuller *et al.* 2005; Garland and Rose 2009). When combined with population-level genome sequencing, such experiments can also help us elucidate the genetic architecture of the selected traits by revealing the quantitative trait loci (QTLs) that underlie observed adaptive responses (Burke *et al.* 2010; Schlötterer *et al.* 2015). This rationale forms the basis of the evolve and resequence (E&R) method for QTL detection (Turner *et al.* 2011; Long *et al.* 2015), in which one seeks to identify the alleles that have systematically changed in frequency over the course of a selection experiment. Such E&R experiments have now been successfully performed in a wide range of study systems (e.g. *Escherichia coli* (Barrick *et al.* 2009; Tenaillon *et al.* 2012), yeast (Parts *et al.* 2011; Lang *et al.* 2013), *Drosophila melanogaster* (Burke *et al.* 2010, Zhou *et al.* 2011, Turner *et al.* 2011), other *Drosophila species* (Seabra *et al.* 2018; Kelly and Hughes 2019), *Caenorhabditis* species (Teotónio *et al.* 2017), and mice (Chan *et al.* 2012; Castro *et al.* 2018)). For example, with E&R experiments, Tenaillon *et al.* (2012) uncovered 600 loci associated with high-temperature tolerance in *E. coli*, and Burke *et al.* (2010) identified several dozen genomic regions responding to selection for accelerated development in *D. melanogaster.*

In higher eukaryotes, practical constraints typically impose severe limits on the size of the experimental population and the number of generations an E&R experiments can be conducted for. Selection pressure is therefore typically kept high so that an adequate trait response can still be achieved. One consequence of such small population size and strong selection is that effective population sizes tend to be low in these experiments, resulting in high levels of genetic drift. In addition, because recombination will be less effective at breaking up linkage in a short experiment, there could be substantial hitchhiking of neutral alleles (Smith and Haigh 1974) as well as Hill-Robertson interference (Hill and Robertson 1966) between selected alleles. All of these factors can limit power and introduce false positives in E&R experiments (Kessner and Novembre 2015).

The first studies to assess the power of E&R experiments for QTL detection in higher eukaryotes used forward-in-time population simulations to model evolutionary dynamics at individual QTLs (Kofler and Schlötterer 2014; Baldwin-Brown *et al.* 2014). These studies provided important insights into how detection power is affected by basic population genetics parameters such as recombination rate, linkage disequilibrium (LD), and the levels of neutral diversity in the regions surrounding the QTLs. In addition, they explored how different aspects of the experimental design such as selection strength, population size, duration of the experiment, and number of replicates can be tuned to maximize detection power.

However, two assumptions of these early studies have turned out to limit the generalizability: First, allele frequency dynamics at individual QTLs were modeled as independent selective sweeps, parameterized by constant selection coefficients. Such models will often fail to capture key aspects of the evolutionary dynamics of QTLs underlying polygenic traits (Burke *et al.* 2010; Kessner and Novembre 2015; Franssen *et al.* 2017). Second, these studies either focused on regions with high recombination (Kofler and Schlötterer 2014) or modeled only a single QTL (Baldwin-Brown *et al.* 2014). This effectively neglects the possibility of Hill-Robertson interference between QTLs, which can affect the evolutionary dynamics in complex ways, thereby impinging on QTL detection power (Hill and Robertson 1966; Smith and Haigh 1974; Lang *et al.* 2013; Kessner and Novembre 2015). Thus, to more accurately describe polygenic trait evolution in E&R experiments, we need to adopt more realistic quantitative genetic models in which the selected trait is defined explicitly and the loci underlying the trait are modeled in the explicit context of a recombining chromosome.

The selection model introduced by Kessner and Novembre (2015) constitutes an important first step in this direction, but it assumed a limited set of genetic trait architectures in which only the number of QTLs was variable. In reality, the traits of interest in E&R experiments could span a considerable variety of genetic architectures, and we typically know very little about this architecture for any given polygenic trait (Hansen 2006; Mackay *et al.* 2009; Gibson 2012). For example, in addition to how many QTLs control a given trait, these QTLs could be distributed uniformly along the chromosome, or they could cluster in certain regions. Effect sizes could be similar among the individual QTLs, or they could vary according to some distribution. The frequencies of the selected alleles might be biased towards lower or towards higher frequencies, as compared to neutrally segregating alleles. The effects of these alleles on the trait might be recessive, dominant, or fall somewhere in between. Furthermore, there could be epistatic interactions of varying complexity among specific sets of QTLs.

Some of these aspects of trait architecture have previously been demonstrated to affect the evolutionary dynamics of trait evolution in E&R experiments. Franssen *et al.* (2017), for example, showed that the effect size distribution and starting frequency of QTLs can profoundly influence their frequency trajectories in response to selection. Similarly, Stetter *et al.* (2018) showed that the effect sizes of QTLs are a key determinant of their frequencies at the end of a selection experiment. As a result, depending on their underlying genetic architecture, certain traits might generally be more suitable for QTL detection in E&R experiments, and for any given trait, there could be systematic biases in terms of which QTLs will be more easily detected and which will be missed. Due to interactions among different factors, this is unlikely to only depend on effect size, so a power analysis considering broader aspects of trait architecture in E&R experiments is required to properly interpret results.

A second limitation in previous studies of detection power in E&R experiments is that they have focused primarily on insect populations like *Drosophila* (Kofler and Schlötterer 2014; Baldwin-Brown *et al.* 2014; Kessner and Novembre 2015), which are well-suited organisms for such experiments due to their short generation times, relative ease at which large populations can be reared, and rich genomic resources. However, for certain questions (e.g. the genomic basis of vertebrate traits), selection experiments on larger and longer-living organisms may be necessary, despite of the additional logistical challenges. E&R experiments on such organisms will typically be restricted to fewer generations, therefore requiring even larger selection intensities to achieve measurable changes in trait value. The greater selection intensity could lead to characteristic differences in evolutionary dynamics compared to experiments carried out over larger numbers of generations and a potential decline in detection power. Nevertheless, many selection experiments have been performed on such larger and longer-living species, even though in many cases their original intentions were not QTL detection (e.g. mice (Oortmerssen and Bakker 1981; Chan *et al.* 2012; Barrett *et al.* 2019), guppies (Houde 1994), silversides (Conover and Munch 2002), voles (Sadowska *et al.* 2008), sticklebacks (Barrett *et al.* 2011), zebrafish (Uusi-Heikkilä *et al.* 2017)). In addition, many common human practices, such as animal domestication and size-selective harvesting (through fishing and hunting), resemble E&R experiments in key aspects such as high selection pressure and specific traits targeted by selection (e.g. domestication of salmonids (Christie *et al.* 2016, Gutierrez *et al.* 2016) and chicken (Rubin *et al.* 2010, Johansson *et al.* 2010, Fallahsharoudi *et al.* 2017), and size-selective harvesting in Atlantic cod (Swain *et al.* 2007, Therkildsen *et al.* 2013) and bighorn sheep (Coltman *et al.* 2003, Pigeon *et al.* 2016)). With high-throughput sequencing becoming cheaper and more widely accessible, genomic data may now be obtained from such experiments and practices. This raises the question of how well time-series data collected over a small number of generations can help us illuminate the molecular basis of selected traits in these larger and longer-living species.

In this paper, we use forward genetic simulations to systematically assess how different aspects of trait architecture are expected to affect the evolutionary dynamics and power to detect QTLs in E&R experiments. Loosely inspired by a size-selection experiment performed on the Atlantic silverside *(Menidia menidia)* to examine impacts of fisheries-induced evolution (Conover and Munch 2002), our model setup is comparable to experimental designs applicable to E&R studies in larger and longer-living species in general (e.g. Sadowska *et al.* 2008; Uusi-Heikkilä *et al.* 2017). The specific aspects of trait architecture that we investigate are the number of QTLs contributing to a selected trait, the clustering of QTLs along the chromosome, the effect size distribution among the QTLs, the starting frequencies of the QTLs, dominance, and epistasis patterns. We show that most of these variables can greatly influence QTL detection power, often in complex ways where the effect of one aspect of the architecture depends on other aspects of the architecture. We also demonstrate that under certain trait architectures, we can use selection experiments spanning as few as four generations to detect a reasonable proportion of QTLs, suggesting that E&R experiments can be a promising approach for studying the genomic basis of adaptation in species with larger size and longer generation time. Motivated by these insights, we further discuss how optimal detection strategies, including detection methods and experimental design, may vary under different quantitative trait architectures.

## METHODS

### Simulation of E&R experiments

We used forward genetic simulations to model E&R experiments in which divergent truncating selection is imposed on a quantitative trait over four consecutive generations. The analysis pipeline consists of the following stages:

1. Burn-in to create genetic variation in a starting population
2. Construction of QTL architecture
3. Selection on the trait
4. QTL detection
5. Power analysis

### Burn-in to generate genetic variation in the starting population

To model a population under mutation-drift equilibrium prior to the selection experiment, we first simulated a 30 Mbp-long chromosome evolving neutrally in a diploid population of *N* = 1,000 individuals for 10N generations. We set the mutation rate to *μ* = 2*10^-8^, corresponding to an expected equilibrium level of nucleotide diversity of *π* = 4*Nμ* = 8*10^-5^. While this value of *π* is comparatively small for many species, it was chosen for computational efficiency and we note that in our analyses of QTL detection power the value of *π* is only expected to affect the absolute number of false positives, but not the false positive rate (i.e. the probability that any given neutral SNP is falsely detected as a QTL, see also Figure S1).

More critical to our power analysis is the rate of recombination, as it will determine the rate at which new allele combinations can arise during the selection experiments and also affect the amount of interference between QTLs and the level of hitchhiking of neutral SNPs with selected alleles. We chose a rate of *r* =1 cM/Mbp for our simulations, which is similar in magnitude to the observed rates in many species of interest (e.g. 0.63 cM/Mbp in mouse (Shifman *et al.* 2006), 0.97 cM/Mbp in dog (Wong *et al.* 2010), 1.5 cM/Mbp in zebra finch (Backström *et al.* 2010), 3.11 cM/Mbp in three-spined stickleback (Roesti *et al.* 2013)). Under these parameters, linkage disequilibrium (LD), as measured by r^2^, decays by 50% over a distance of approximately 50 kbp in our simulated population.

### Assignment of QTLs and construction of the trait architecture

In our standard trait model, we randomly selected *n* of the existing neutral SNPs after the burn-in to become the QTLs affecting the trait. We then randomly picked half of these SNPs to which we assigned a positive effect (+1) to the derived allele and a zero effect to the ancestral allele. For the other half of these selected SNPs, we assigned a negative effect (−1) to the derived allele and a zero effect to the ancestral allele. We assumed additive dominance relationship (h = 1/2) at individual QTLs and also additive effects across QTLs. Under this model, the average trait value in the population is expected to be zero at the start of a selection experiment and the distribution of trait values among individuals should be approximately Gaussian (as long as *n* is sufficiently large), consistent with many polygenic quantitative traits in nature (Mackay 2009). We did not model the effect of environmental factors on trait value (i.e. the broad-sense heritability of the trait is set to a value of one in our simulations).

Six aspects of this standard model were varied to explore different trait architectures: (i) the number of QTLs, (ii) the clustering of QTLs along the chromosome, (iii) the effect size distribution of QTLs, (iv) the initial allele frequency distribution of the SNPs chosen to become QTLs, (v) the dominance relationship between the ancestral and derived alleles, and (vi) the presence of epistatic interactions among pairs of QTLs. The specific implementations of each of these architectures are described in the relevant sections below.

### Selection experiment on the trait

In the selection experiment, the population is subjected to divergent truncating selection, generating two separate lines from the burn-in population: a high-trait-value line and a low-trait-value line. For the “high” line, we selected the 10% of individuals with the highest trait values in every generation to become the parents for the next generation (obtained by Wright-Fisher sampling). The “low” line was generated analogously by choosing the 10% of individuals with the lowest trait value as parents for the next generation. Population size was kept fixed at 1000 individuals per generation in each line, and each line was run for four generations of truncating selection following the design of the silverside experiment that served as a motivating example for this study (Conover and Munch 2002). Because of this short duration, the impact of new mutations occurring during the selection experiment should be negligible, and we therefore set the mutation rate to zero after the burn-in while recombination events continue to occur at a rate of 1 cM/Mb.

### QTL detection

In each generation, we took a random sample of 50 individuals and measured the allele frequencies of all SNPs in the sample. Following Kessner and Novembre (2015), we took the absolute values of sampled allele frequency differences between the last generations in the “high” and “low” lines at each SNP (denoted by D) as a summary statistic for QTL detection.

In addition to this simple and intuitive D-value, several more sophisticated, model-based detection methods have been developed in recent years that can take advantage of the full allele frequency trajectory estimated across subsequent time points (Malaspinas 2016). After a comprehensive literature review, we selected two representatives of such methods for comparison with the simple D-value: WFABC (Foll *et al.* 2015) and ApproxWF (Ferrer-Admetlla *et al.* 2016). Both of these methods are based on a classic selective sweep model parameterized by fixed selection coefficients. WFABC employs an approximate Bayesian computation framework, in which a large number of simulations are compared to identify the simulated datasets that are most similar to the actual data and compute posterior probabilities of selection coefficients (Foll *et al.* 2015). ApproxWF uses a “mean transition time approximation” to discretize the continuous diffusion process and infers the selection coefficient via a Bayesian approach (Ferrer-Admetlla *et al.* 2016).

After a series of tests, we adjusted some parameter values in WFABC from the default to optimize its detection power on our standard model, by increasing the number of simulated datasets to 1,000,000, lowering the acceptance rate to 0.00, and assigning a uniform prior to selection coefficients with an upper bound of 1 and lower bound of −1. With ApproxWF, we used its default Markov Chain Monte Carlo settings, with 10,000 iterations and 51 frequency states distributed on quadratic grid, and assigned a normal prior to selection coefficients, with mean of 0 and standard deviation of 0.1, truncated at 1 and −1. We also set a fixed dominance coefficient of 0.5 to reduce the computational complexity in ApproxWF.

Both methods output a posterior distribution for the selection coefficient for each SNP, and from this distribution, we calculated a mean selection coefficient s for each SNP, as well as a posterior probability *p* for s>0, as recommended by authors of these methods (Foll et al. 2015, Ferrer-Admetlla et al. 2016). We applied WFABC and ApproxWF to the high-trait-value line and the low-trait-value line separately, and then took the average of s and *p* across the two lines while accounting for the directionality of selection (i.e. 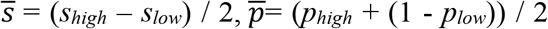). Lastly, since +1 alleles and −1 alleles are not distinguished and both are considered as QTL in the power analysis, the signs of selection coefficients would not matter, so we took 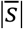 and 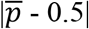 as the final summary statistics for each SNP. After testing for the detection power of these statistics, we chose to use 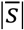 for WFABC and 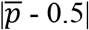 for ApproxWF to maximize their performance.

### Power analysis

We calculated receiver operating characteristic (ROC) curves to evaluate QTL detection performance using 100 simulation replicates for each scenario. False positive rates were defined as the percentage of neutral SNPs identified as QTLs for a given signal threshold. To evaluate power (the true positive rate) for a given signal threshold, we deployed two different methods: The first method simply measures the proportion of QTLs correctly identified. In simulations where effect sizes were not equal among QTLs, we also used a simple variation of this method, where we weighted QTLs by their effect sizes. The second method follows Kessner and Novembre (2015) and measures the proportion of genetic variance in the first generation explained by the detected QTLs, thereby also taking variation in the allele frequencies into account. This method would give less weight to the detection of a low-frequency QTL compared with an intermediate-frequency QTL, because the latter would have contributed more to the initially present trait-variance in the population. Which method is more appropriate in practice depends on the specific objective of the experiment: if the goal is to identify QTLs that are important to the trait regardless of their prevalence in nature, the first method should be chosen, whereas the second method should be chosen if the goal is to identify those QTLs that are most important for explaining trait-variance in the initial population. We will only report the result from the first method unless the two methods generate qualitatively different results, in which case we will present both.

To produce an individual ROC curve, we first specified 100 evenly-spaced signal thresholds spanning the range of observed per-SNP values of the given summary statistics (*D*, 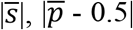) among all replicate runs. For each threshold value in each replicate run, we categorized SNPs with summary statistics exceeding that threshold value as being detected as a QTL in that run. We then took the mean power and false positive rate across all replicate runs to add one point to the ROC curve. This process was reiterated for each threshold value.

### Data availability

All simulations were conducted within the individual-based forward genetic simulation framework SLiM 2.4.1 (Haller and Messer 2016). Two SLiM scripts were created for our analyses, one for the burn-in process and the other for the selection experiment. Two shell script can be used to run each of these SLiM scripts in a command-line environment, so that repeated simulations can be automated on either a local machine or a remote server. Parameter values are also defined through these shell scripts and users can easily edit them to implement custom simulation scenarios. All data analyses and visualizations are implemented using R, with packages “tidyverse” (Wickham and RStudio 2017), “data.table” (Dowle *et al.* 2019), and “cowplot” (Wilke and RStudio 2019). The SLiM and shell scripts for the simulation are available in supplementary materials and online at https://github.com/MesserLab/evolve-resequence-simulation. The data used in this paper can easily be reproduced using these scripts and the scripts can also be customized for other test cases. The R scripts for data analyses and visualization are also available in the same GitHub repository.

## RESULTS

### Selection response in the standard QTL model

Figure 1A shows the change in average trait value (estimated across all individuals in the population) for the “high” and “low” lines in 100 simulated experiments under our standard model with *n* = 100 QTLs (Figure S2 shows results for a model with *n* = 10 QTLs). Average trait values can be seen to change consistently in the selected direction, while genetic variance generally declines with diminishing returns over the course of a single experiment (Figure 1B). Note, however, that the theoretical maximum/minimum trait value in our standard QTL model would be +/-100 had all of the +1 or all of the −1 alleles, respectively, fixed in a population. The maximum trait values achieved in our simulations were typically less than a third of these maximum values. This is primarily due to the fact that many low-frequency alleles at QTLs are lost due to drift or interference between neighboring QTLs even when they should have been favored by selection. Overall, these results demonstrate an effective selection response in our model that is consistent among replicates, suggesting that we should be able to observe meaningful evolutionary dynamics in our experimental setup.

**Figure 1.**
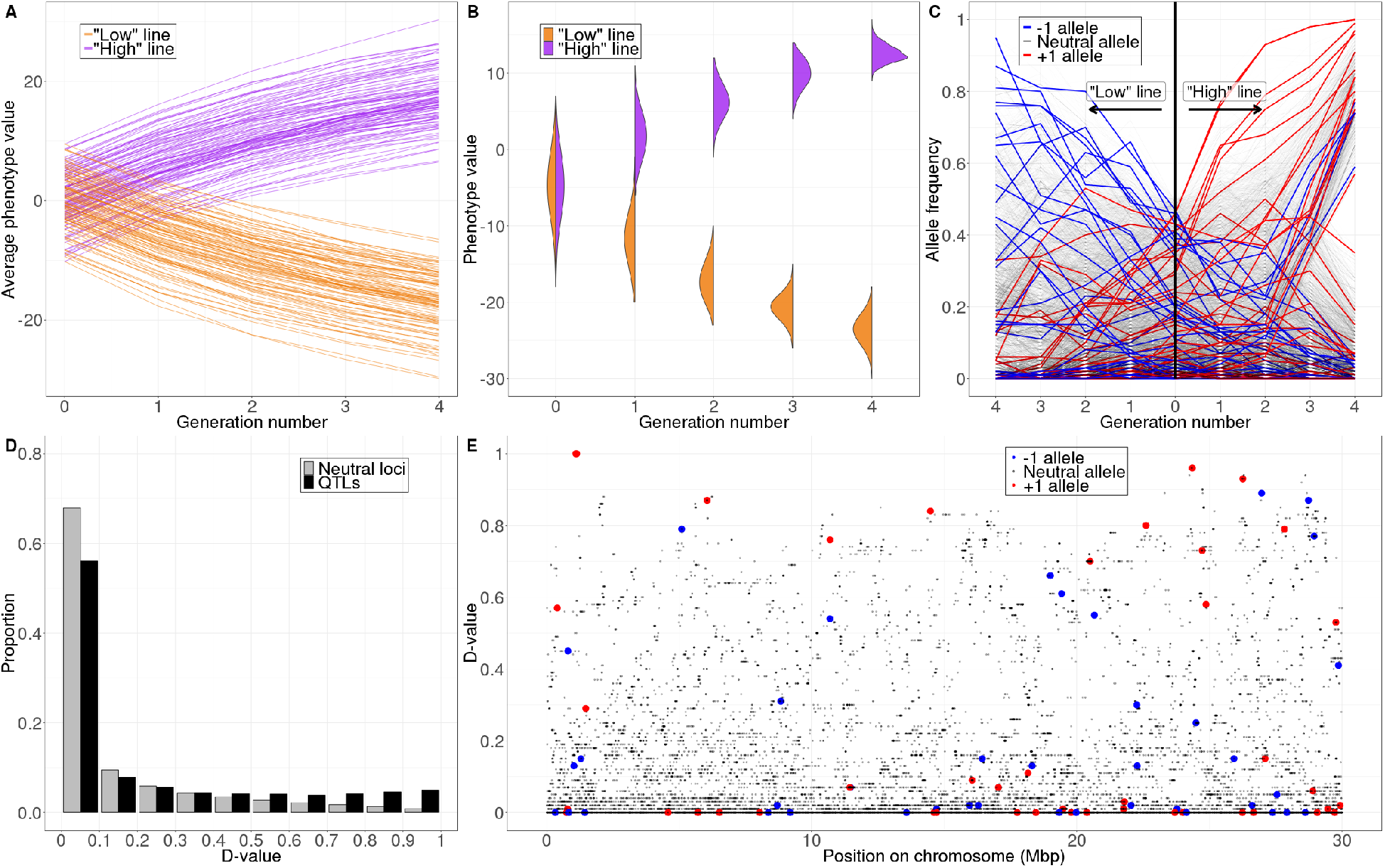
Simulation result under our standard model with 100 QTLs. (A) Change in the average phenotype values in all 100 simulation replicates. Each line represents a selection line in one simulation replicate. (B) Change in the distribution of trait values in the population over one single simulation run. (C) Change in sampled minor allele frequencies at neutral loci and QTLs in one single simulation run. The left half of the figure shows the “low” line and the right half shows the “high” line. (D) Distribution of *D*-values per locus across all 100 simulation replicates grouped by neutral loci (grey bars) vs QTLs (black bars). (E) *D*-values of neutral loci and QTLs along the simulated chromosome in one single simulation run.

Figure 1C shows sampled allele frequency trajectories at all SNPs in the “high” and “low” lines from a single experiment. As expected, minor alleles at the individual QTLs tend to change in frequency in the selected direction (i.e., in the “high” line the +1 alleles tend to increase while the −1 alleles tend to decrease in frequency, and vice versa in the “low” line). However, several complexities of the allele frequency dynamics are revealed in this figure: First, many alleles that should have been favored by selection did not actually rise consistently in frequency. This is particularly common for alleles that start at low frequency, which often get lost in both selection lines. But even after favored alleles reach intermediate frequency, they can still subsequently drop in frequency due to interference with other linked QTLs. Similarly, alleles that are disfavored by selection can hitchhike to higher frequencies when they are located on haplotypes with a net excess of favored alleles. These linkage effects produce dynamics that are quite different from a model of independent selective sweeps.

Figure 1D shows the distribution of D-values across SNPs over 100 simulation runs of our standard model with *n* = 100 QTLs. The distribution is heavily peaked at D-values equal or close to zero for both neutral and QTL SNPs, which is expected because the derived alleles at most SNPs will be at low frequency at the start of the experiment, and thus prone to being lost to drift in both the “high” and “low” lines. This will generally limit detection power when measured as the overall fraction of QTLs identified, given that many of the initially present alleles at QTLs will be lost in an experiment. However, we also see that among those SNPs with high D-values, functional SNPs are strongly enriched over neutral ones, suggesting that *D*-value should have some power in detecting QTLs under this experimental setup. Figure 1E shows the distribution of D-values along the chromosome in one simulation run with 100 QTLs, demonstrating that neutral SNPs with high *D*-values are not necessarily always close to the QTLs, but can be found across the whole chromosome.

### Performance of different detection methods under the standard QTL model

We first compared the performance of D, WFABC, and ApproxWF to detect the QTLs in our standard model, assuming a trait comprised of 10 QTLs with equal effect size. Even though SNP density is comparatively low in our standard model (~14,000 SNPs along the 30 Mbp chromosome), the runtime of the two model-based methods still exceeded our practical limits, so we further reduced the nucleotide diversity by a factor of 10 in these simulations (for this comparison only). We then tested the model-based methods supplying them either with the entire allele frequency trajectory (i.e. allele frequency estimates at all five time-points of the experiment), or only the allele frequencies at the beginning and the end of the experiment.

Surprisingly, WFABC and ApproxWF typically had lower power to correctly detect QTLs (i.e. lower true detection rate for a given false negative detection rate) than the simple D-value, even when they were provided the full allele frequency trajectory at all five time points (Figure 2). This may in part be due to the fact that these methods were not explicitly developed for a divergent selection scenario resulting in two opposingly selected lines. However, when we restricted the experiment to only one direction of selection, *D* still performed similarly and often better than WFABC and ApproxWF (Figure S3). One possible explanation for the poorer performance of the model-based approaches is that the allele frequency dynamics at QTLs in our truncating selection scenario may not typically follow the classic sweep model these methods assume, as has already been observed in previous studies (Burke *et al.* 2010; Kessner and Novembre 2015; Franssen *et al.* 2017). Consequently, we decided to use only the D-value for all further analyses in this study.

**Figure 2.**
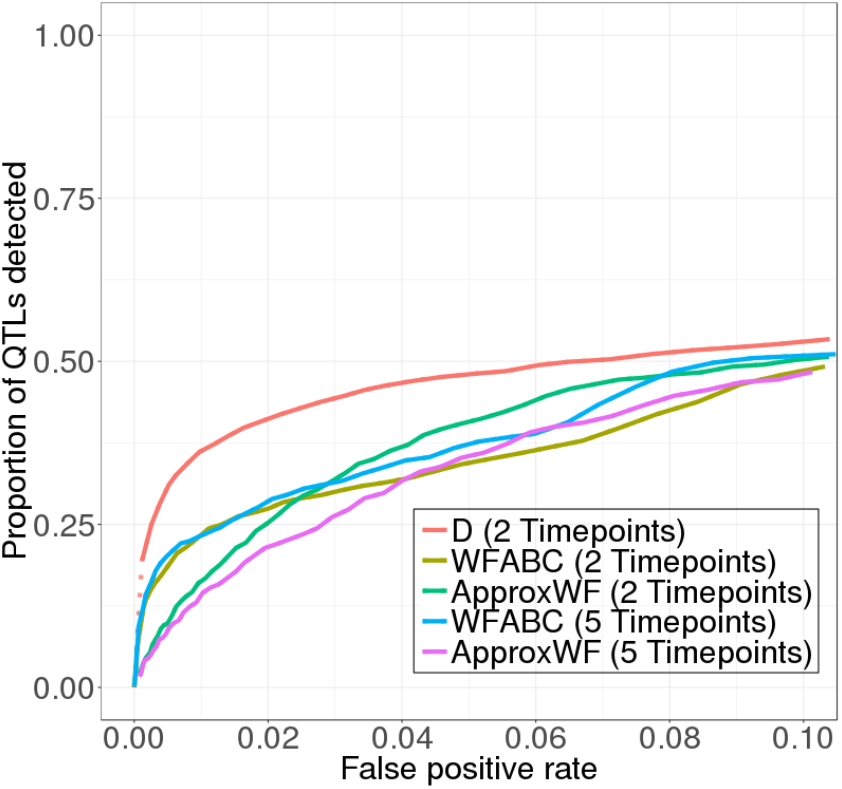
Detection power varies across detection methods used. Having the entire allele frequency trajectory slightly improves the power of WFABC but not ApproxWF, although *D* has the highest detection power regardless. This comparison used the standard model with 10 QTL and 10-times reduced nucleotide diversity due to the long run-time of the two model-based detection methods.

### Effects of trait architecture on detection power

#### Number of QTLs affecting a trait

To test how the number of QTLs contributing to a trait affects detection power, we systematically varied the number of SNPs assigned to be QTLs in our simulations. Figure 3 shows a comparison of the ROC curves among models with 2, 10, 20, 50, 100, and 200 QTLs, while all other aspects of the model were kept the same as in the standard model. Consistent with previous results (Kessner and Novembre 2015), we find that larger numbers of QTLs generally resulted in lower detection power, presumably due to increased interference between QTLs. As the number of QTLs increases, individual QTLs will tend to be located closer to each other, decreasing recombination rate between them. Recombination will then be less effective at creating “optimal” haplotypes that carry a large number of favored but only few unfavored alleles. A complementary effect is that more QTLs also mean less effective selection on every single one of them, because the relative contribution of each individual mutation to the overall variance in trait value gets smaller (Barton and Turelli 1989).

**Figure 3.**
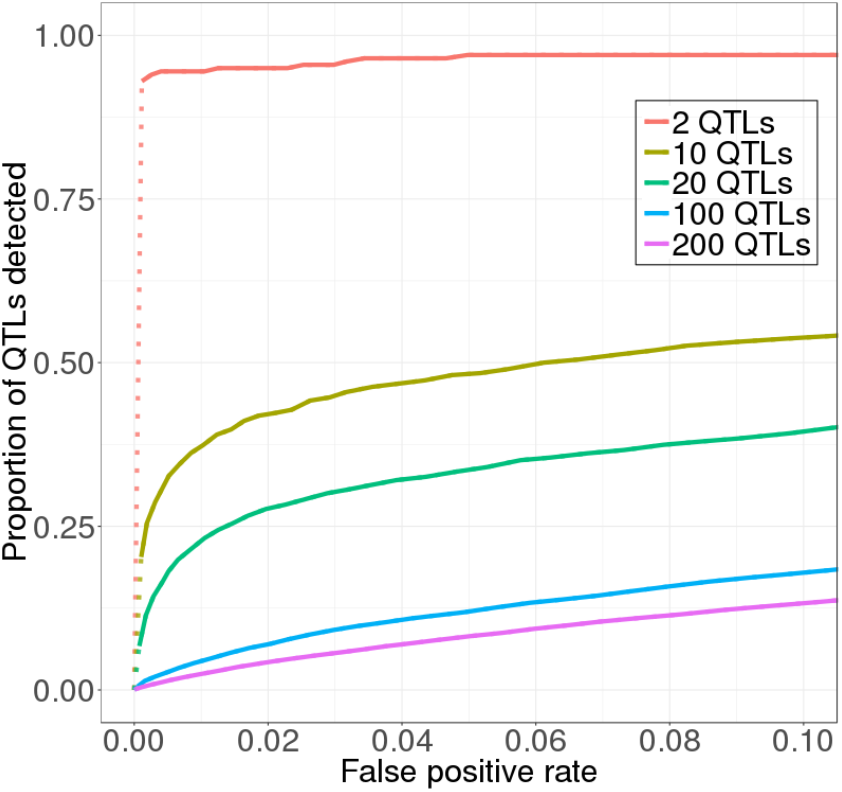
Detection power is lower when more QTLs underlie the trait under selection.

Note that the overall detection power is rather low in our standard model with 100 QTLs. In that case, we detected only ~13% of the QTLs at a false positive rate of 0.05 (i.e. 5% of the neutral loci are falsely identified as QTLs). This is because most QTLs start out at low frequency and have a high probability of getting lost over the course of the experiment. However, since intermediate-frequency QTLs are more likely to be detected, when weighting QTLs by their contribution to genetic variance in the first generation (which should be much higher for intermediate-frequency than low-frequency QTLs), power improves significantly (Figure S4). For example, at the same false positive rate of 0.05, the detected QTLs were responsible for more than 40% of the genetic variance present in the first generation (Figure S4).

#### QTL clustering

In our standard model the QTLs are positioned uniformly along the chromosome, as we chose them randomly from preexisting SNPs. However, for some traits, QTLs could cluster along the chromosome. This is frequently observed among domestication-related traits in crops, for example (Cai and Morishima 2002; Burger *et al.* 2008). To test how such clustering affects detection power, we compared our standard QTL model with a model in which all QTLs were drawn from only those SNPs that were located within a much shorter genomic sub-region 3 Mbp in length, located at the center of the chromosome.

For a model with only 10 QTLs, we found that such clustering lowers detection power compared to a more uniform distribution (Figure 4A & S5A). This is likely due to increased linkage between QTLs, resulting in an effectively lower rate at which recombination can create haplotypes with a large number of favored alleles. Clustering therefore has a similar effect as increasing the overall number of QTLs while keeping the length of the chromosome constant.

**Figure 4.**
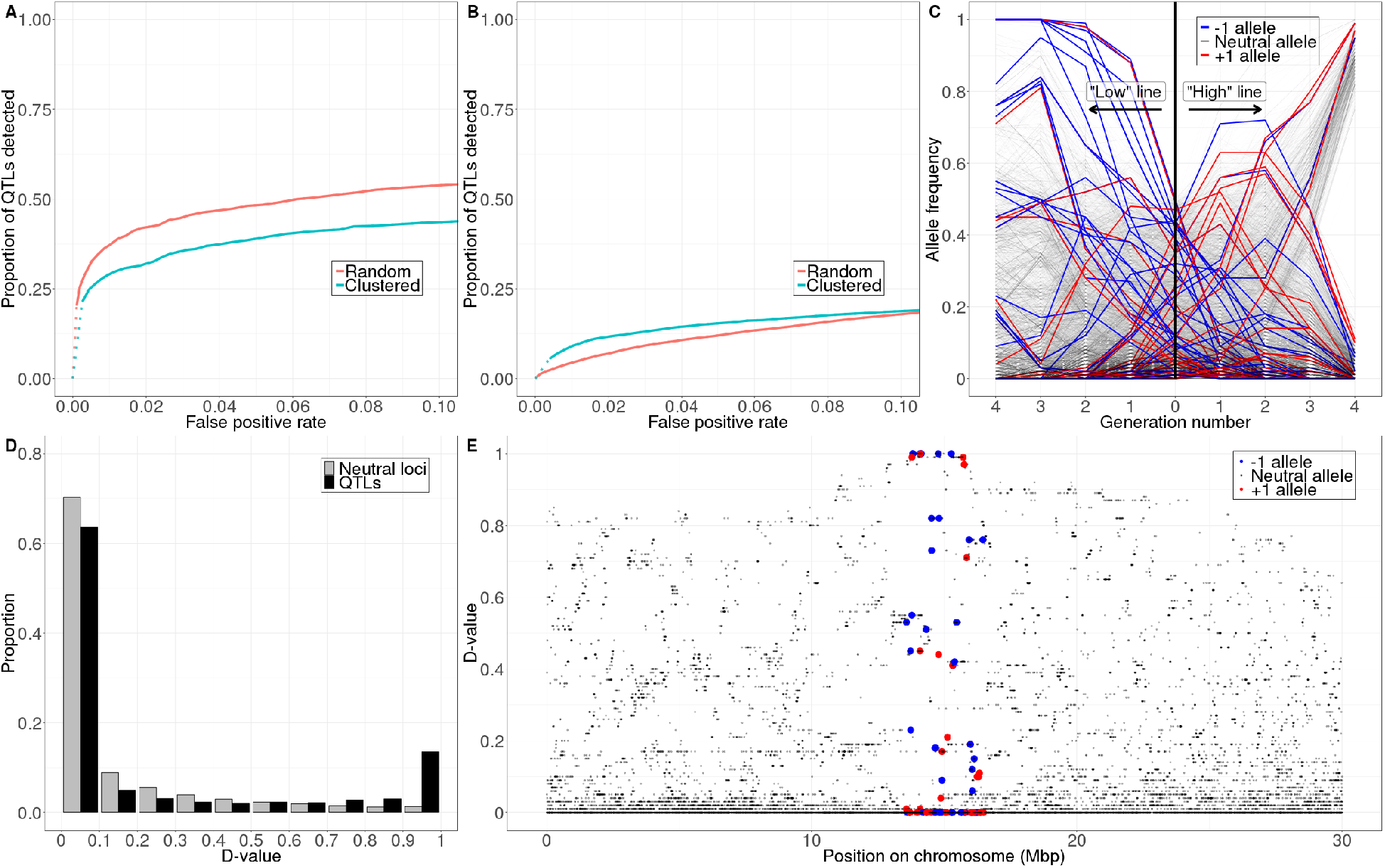
The clustering of QTLs has different effects on detection power depending on the number of QTLs underlying the trait under selection. (A) With 10 QTLs, clustering reduces detection power. (B) With 100 QTLs, clustering increases detection power except for at very high false positive rates (>0.1). (C) An example of allele frequency trajectories when 100 QTLs are clustered in a small region on the chromosome. Note that many SNPs on the same haplotype quickly sweep to fixation in tandem in the “low” line, and see Figure 1C for comparison with the standard model. (D) Such haplotype sweeping results in more QTLs with extreme *D* values (see Figure 1D for comparison with the standard model). (E) The distribution of *D*-values along the chromosome after a haplotype sweeping (see Figure 1E for comparison with the standard model).

However, this behavior becomes more complicated as the number of QTLs increases further. In a model with 100 QTLs, we found that the effect of clustering on detection power depends on what false positive rate is deemed tolerable. For false positive rates below 0.1, clustering actually increases detection power (Figure 4B & S5B). A possible explanation for this is that when there are many QTLs clustered within a short region on the genome, it is more likely that already in the beginning of the experiment a short haplotype exists on which many alleles with effects of the same direction are co-located. Unlikely to be broken by recombination, such a haplotype will be able to quickly sweep to fixation, giving a very clear signal of being under selection (Figure 4C, 4D & 4E). Therefore, at a low false positive rate, the scenario where 100 QTLs are clustered on the chromosome tends to have higher power than our standard model.

Furthermore, we note that clustering has a similar effect as decreasing the recombination rate in our simulation. Thus, our results contradict that of Kessner and Novembre (2015), who concluded that increasing recombination always increases detection power. However, our results are consistent with a rich body of literature which shows that increased recombination does not always lead to higher rate of adaptation because it can also destroy “good” haplotypes that are initially present (Slatkin 1975; Charlesworth and Charlesworth 1979; Kirkpatrick 2006).

#### Effect size distribution

Our standard model assumes equal effect sizes of derived alleles of either +1 or −1 at all individual QTLs. We chose this simplistic model because little is known about the actual effect size distributions of complex traits in most biological systems and thus there is not a single ideal distribution. Also, results on other aspects of the trait architecture will be more difficult to interpret if we add an additional layer of stochasticity to the simulations by randomly assigning uneven effect sizes. In reality, however, effect sizes will typically vary among QTLs, and one commonly used model for this is an exponential distribution for effect sizes (Orr 1998; Otto and Jones 2000).

Figure 5 shows how QTL detection power is affected when effect sizes in our standard model are no longer assigned equal values, but instead are drawn from an exponential distribution with means +1 or −1, respectively. Whether this increases or decreases detection power depends on how we define our measure of power. If power is defined simply as the proportion of QTLs detected regardless of their effect sizes, power is lower in the model with exponentially-distributed effect sizes compared with our standard model assuming constant effects (Figure 5A & 5D). One possible explanation for this is that in the exponentially-distributed model, there will be many QTLs with small effect that are practically neutral and are likely to get lost due to drift or interference. This means that there would be a smaller proportion of “effective” QTLs that can still be detected overall.

**Figure 5.**
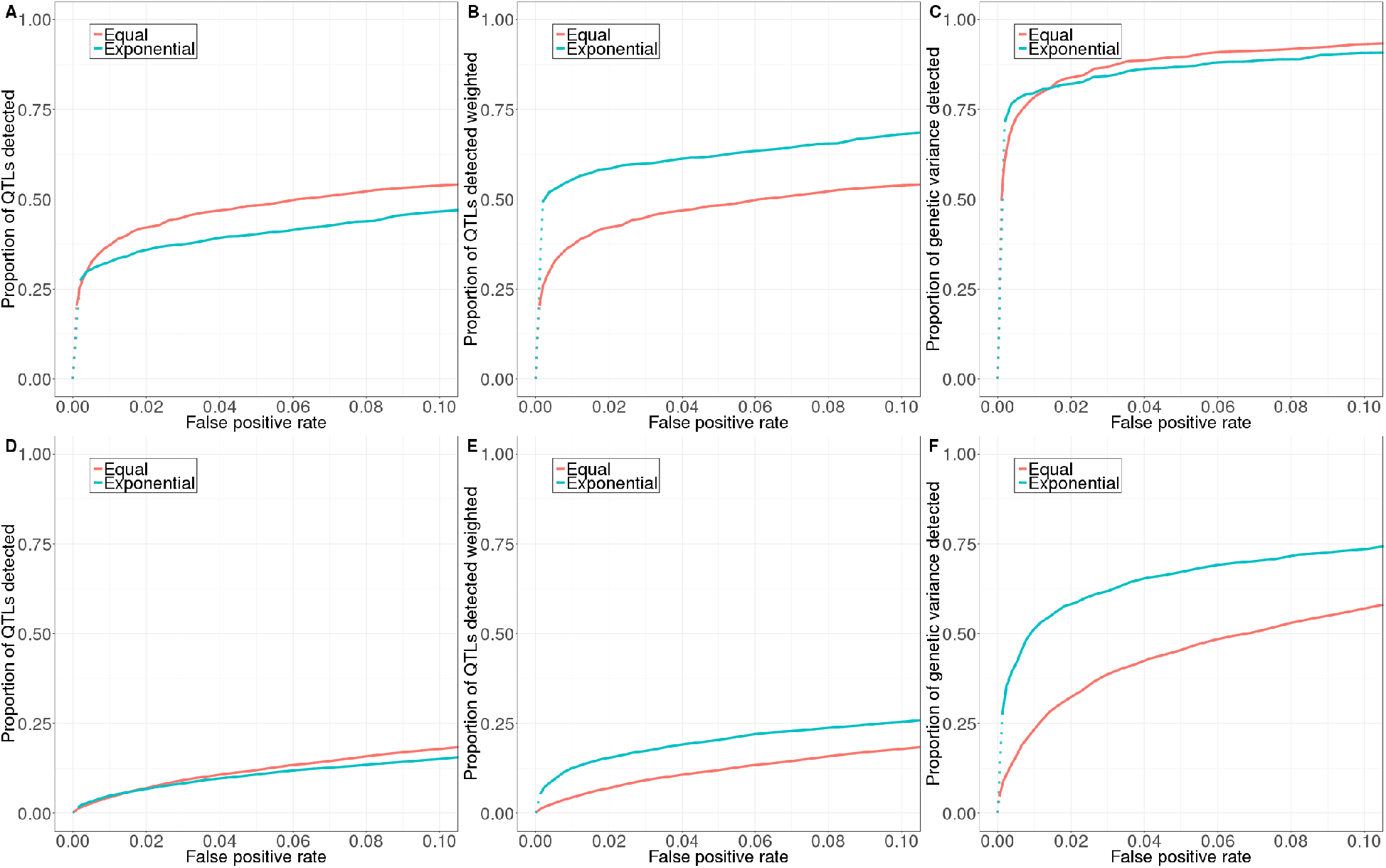
An exponential distribution of effect size affects detection power differently depending on how power is evaluated and how many QTLs underlie the selected trait. All comparisons are conducted with the standard model of equal effect sizes. **Top**: 10 QTLs. **Bottom**: 100 QTLs. (A) (D) Exponential distribution of effect size decreases power when measured by the proportion of QTLs detected. (B) (E) By contrast, exponential distribution of effect size increases power when measured by the proportion of QTLs detected, weighted by their effect sizes. (C) (F) When power is measured by the proportion of genetic variance explained, an exponential distribution of effect size yields similar power when there are 10 QTLs but increases power when there are 100 QTLs.

However, when we weight individual QTLs by their effect sizes, these small-effect QTLs will contribute minimally to power, whereas the few large-effect QTLs become much more important (Chevalet 1994). Similar to the effect of lowering the number of QTLs as discussed above, these few large-effect QTLs will interfere less with each other because they are less densely distributed on the chromosome and are more likely to be detected. As a result, when QTLs are weighted by their effect sizes, the model with exponentially distributed effect sizes yields higher power than the constant effect size model (Figure 5B & 5E).

When we measure the proportion of genetic variance explained by detected QTLs in the first generation, exponentially distributed effect sizes have two opposing effects. On the one hand, as discussed above, large-effect QTLs are more likely to be detected, and they contribute more to the genetic variance given the same starting frequency. On the other hand, large-effect QTLs are less likely to get lost due to drift or interference even when they start at low frequency, so intermediate-frequency QTLs are less overrepresented among detected QTLs (Figure S6), which would then decrease the proportion of initial genetic variance explained by the detected QTLs compared to our standard model. When there are fewer QTLs, the first effect is weaker than the second (Figure S6A & S6B), so detection power becomes similar between the standard and exponential models (Figure 5C). When there are more QTLs, there is also more interference among large-effect QTLs and thus the second effect becomes weaker (Figure S6C & S6D), resulting in higher power for the exponential model (Figure 5F).

#### Allele frequency distribution

We still know very little about what an appropriate distribution for the starting allele frequencies at the QTLs of any given trait in our experiment would be. This distribution could differ markedly among traits and will also depend on the evolutionary history of the population. For example, a trait that has been under strong stabilizing selection for a long period of time would likely have a very different distribution of allele frequencies at its QTLs compared with a trait that has recently experienced a change in trait optimum, or a trait whose optima would have varied in different parts of the population. In our standard QTL model, we have simply assumed that allele frequencies at QTLs resemble those of neutral SNPs.

Now, instead of randomly selecting SNPs, we only selected SNPs with minor allele frequencies above or below a certain cutoff to become QTLs in the simulation. When we restricted our selection to SNPs with minor alleles at a frequency lower than 5%, detection power decreased consistently regardless of the method to evaluate power (Figure 6). This is presumably because these alleles are more likely to get lost due to drift or interference regardless of the direction of selection.

**Figure 6.**
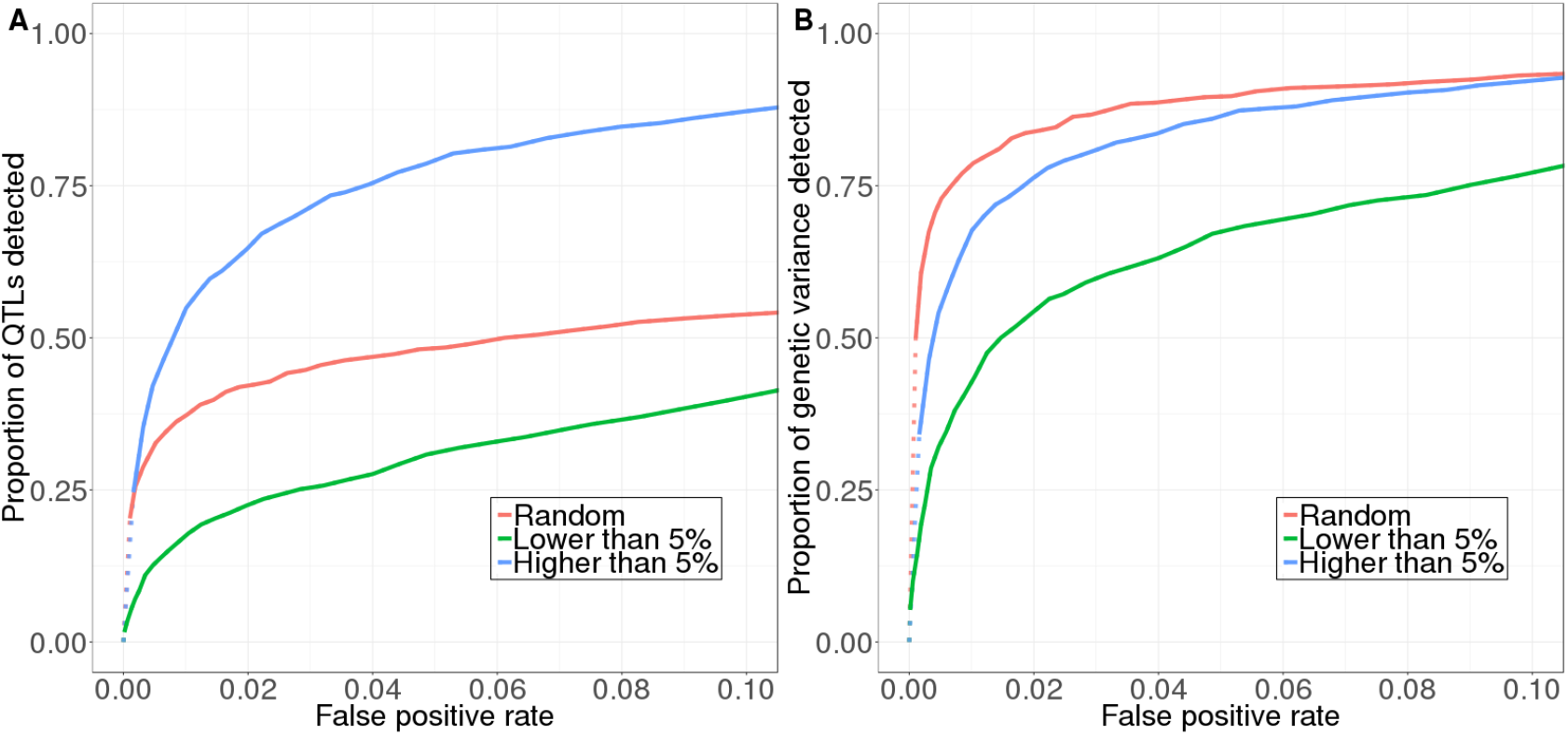
Starting frequency of minor alleles affects power differently depending on how power is evaluated. (A) Power evaluated as proportion of QTLs detected. (B) Power evaluated as proportion of genetic variance explained by detected QTLs. Models with 10 QTLs are used in this figure.

Conversely, when we restricted our selection to only those SNPs with minor alleles at a frequency higher than 5%, these alleles are less likely to get lost. When they are favored by selection, they are also more likely to be recombined together with other favored alleles. This is consistent with the observation that a higher proportion of total QTLs can be detected in such a scenario (Figure 6A). However, the result is different when we evaluate power by the proportion of the starting genetic variance (Figure 6B). The reason is that in the standard model, QTLs that start at intermediate frequencies are more likely to be detected and their detection can explain a higher proportion of starting genetic variance (Figure S7A). When all QTLs start at more intermediate frequencies, they all explain a similar proportion of the starting genetic variance, and the likelihood of them being detected becomes more independent of their starting frequencies (Figure S7B), so even though more QTLs can be detected, the proportion of initial genetic variance explained ends up lower.

#### Dominance

Previous QTL models have typically assumed additive effects between the two alleles at individual QTLs, which is what we also adopted in our standard model. However, dominance effects at QTLs, as well epistatic effects among QTLs, could play an important role in many traits (Shao *et al.* 2008; Mackay 2014; Chen *et al.* 2015).

We first tested how dominance relationships affect QTLs detection power in our model. Given how our QTLs are initially assigned, derived alleles tend to be the minor alleles in our model. When these derived alleles are completely dominant, heterozygotes will exhibit the same phenotype as derived homozygotes, and selection thus cannot distinguish between them. As a result, positively-selected derived alleles tend to first increase in frequency, but then accumulate at intermediate frequencies. This produces a less conspicuous signal of selection than for alleles that reach higher frequencies over the course of the experiment (Figure 7C & 7D). At low false positive rates, detection power is therefore lower in this scenario than in the standard scenario where alleles are assumed to be additive (Figure 7A & 7B). At a higher false positive rate, however, a crossover pattern in the power curves is observed (Figure 7A & 7B), since lower frequency alleles are less likely to be lost due to interference than in the standard model.

**Figure 7.**
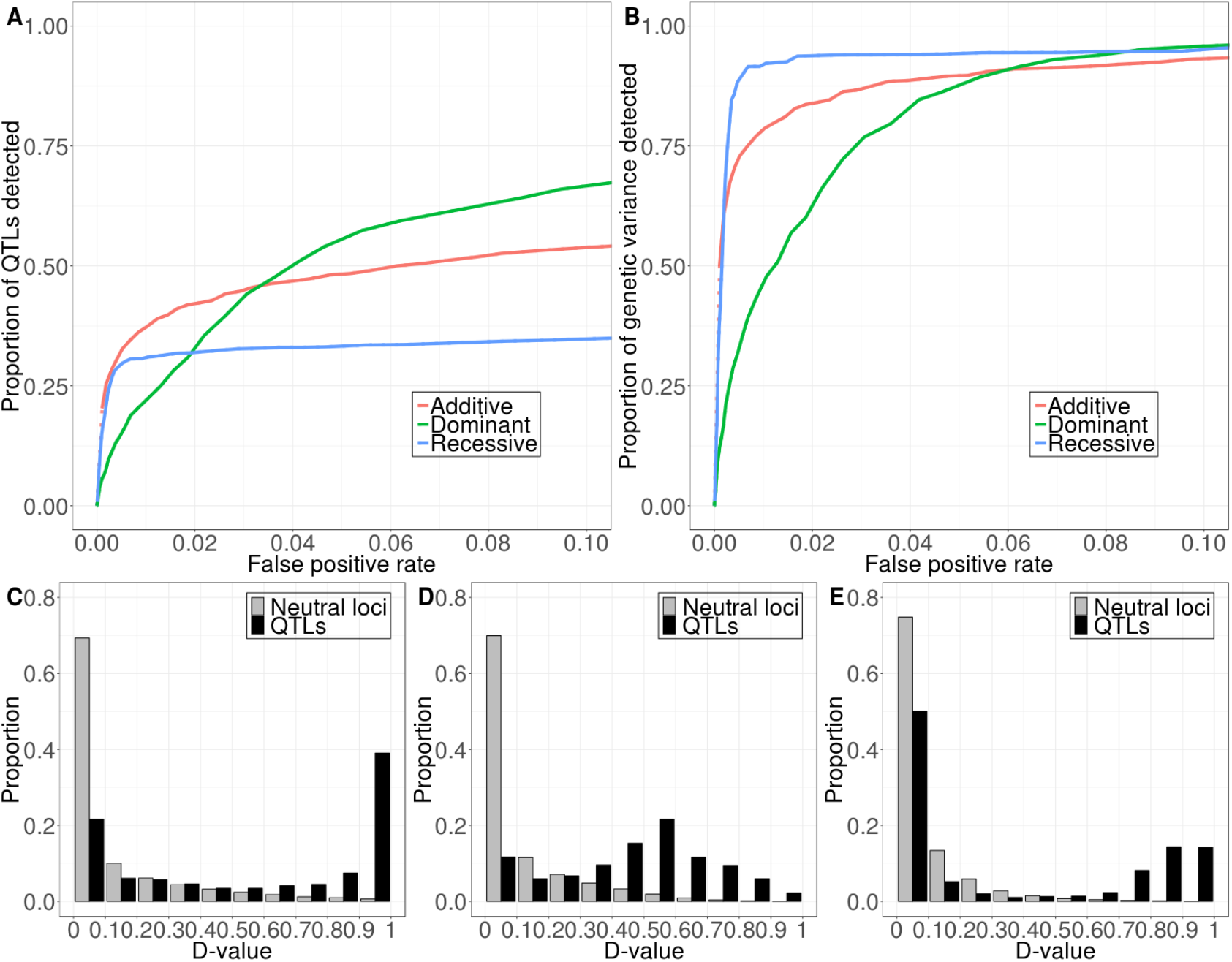
Dominance affects power differently depending on how power is evaluated and what level of false positive rate is tolerated. **Top**: ROC curves (A) Power evaluated as proportion of QTLs detected. (B) Power evaluated as proportion of genetic variance explained by detected QTLs. **Bottom**: Distribution of D-value over 100 simulation replicates (C) Additive. (D) Dominant. (E) Recessive. Models with 10 QTLs are used in this figure. Additive: heterozygotes express intermediate phenotype. Dominant: heterozygotes express the same phenotype as derived homozygotes. Recessive: heterozygotes express the same phenotype as ancestral homozygotes.

When derived alleles are completely recessive, only those with high starting frequency can form homozygotes and thus be visible to selection (Figure S8). All other derived alleles behave essentially like neutral alleles initially (Figure 7C & 7E). As a result, an overall lower number of QTLs can be detected (Figure 7A). However, these high-frequency QTLs are also the ones that contribute most to genetic variance in the first generation. Therefore, the proportion of genetic variance detected is even higher than in the standard model (Figure 7B). These results demonstrate again how a relatively simple aspect of the trait architecture – here the dominance relationship at individual QTLs – can affect detection power in complex ways, where the direction of the effect depends on the false positive rate and the definition being used to measure power.

#### Epistasis

We next tested how epistatic interaction among QTLs can affect detection power. For simplicity, we only considered pairwise epistasis here. We further restricted our analyses to additive-by-additive epistasis (i.e. the effects of the two alleles at an individual QTL are always additive if genotypes at other QTLs are fixed). This helps us avoid the potentially confounding effect of dominance. Under these assumptions, we tested the effect of epistasis with our 10 QTLs model, where we randomly selected five epistatic QTL-pairs in each simulation. We explored separately the effect of four major types of epistasis, including synergistic, antagonistic, sign, and reciprocal sign epistasis. Within each type, we further created a “weak” and a “strong” scenario, based on the level of deviation from the non-epistatic model (Table S1). To evaluate power for a given epistasis scenario, we simply measured the overall proportion of the 10 QTLs we detected, since the effect of each individual QTL is difficult to quantify.

Our simulations show that pairwise epistasis generally decreases the power in QTL detection, as would be expected (Figure 8A). However, a lot of variability exists among different epistasis scenarios. Within each type of epistasis, the more it deviates from the non-epistatic case, the less detection power is generally attained. Among different types of epistasis, synergistic epistasis tends to have higher power, while sign and reciprocal sign epistasis tend to have lower power. Since a major effect of epistasis is introducing epistatic genetic variance and lowering the trait’s narrow-sense heritability (*h*^2^), we examined how heritability varied across our epistasis scenarios, and found that heritability is a good predictor of power (Figure 8B). Scenarios that create lower heritability (or higher epistatic genetic variance) tend to have lower QTL detection power. However, sign epistasis is an exception to this rule. It has relatively high heritability throughout the experiment, but its detection power remained low in our simulations.

**Figure 8.**
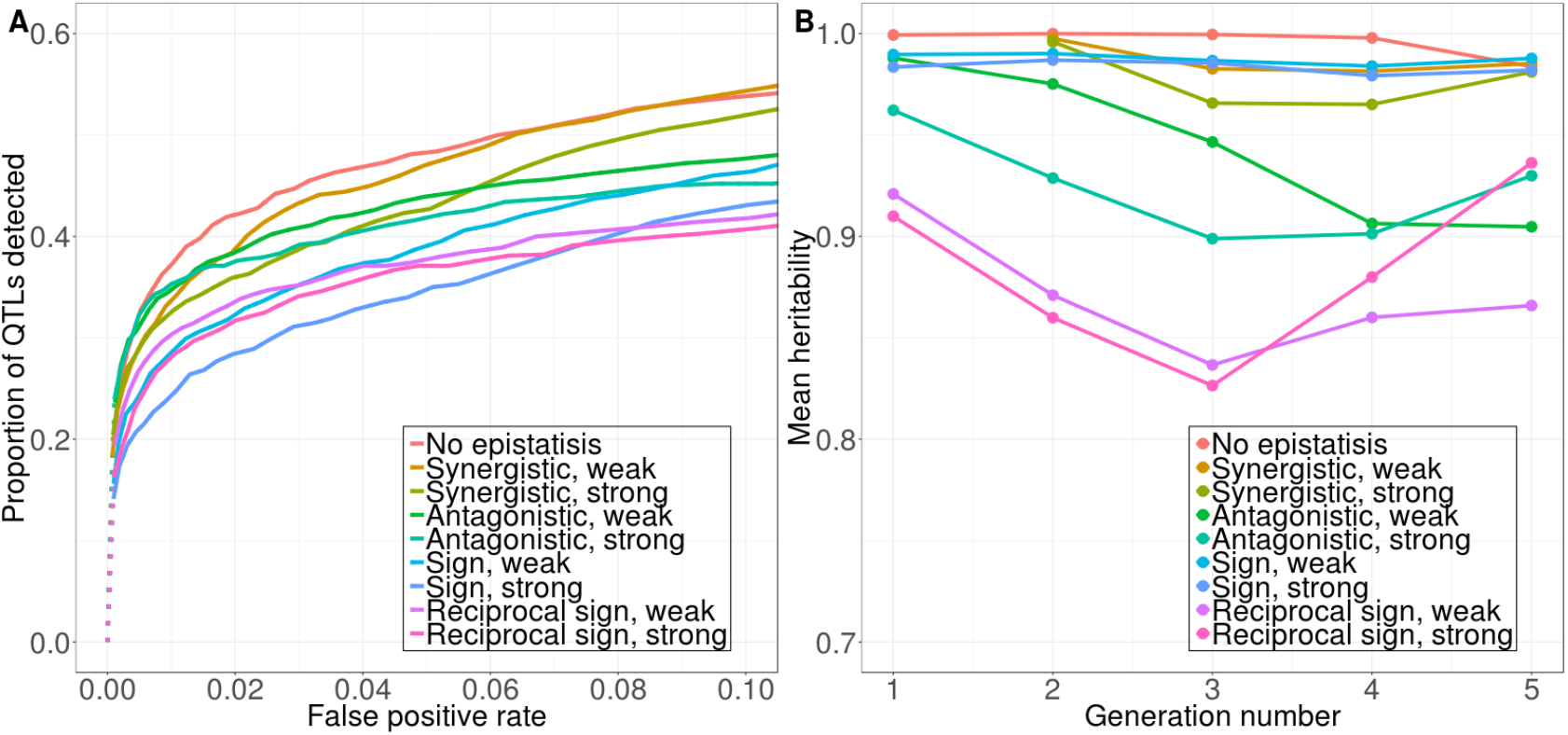
(A) Pairwise epistasis always decreases QTL detection power, but this effect is a lot stronger in certain epistatic scenarios than others. (B) Estimates of narrow-sense heritability for a given epistasis scenario at a given time-point are calculated from the breeder’s equation (Lush 1943), averaged across all 100 simulation replicates.

## DISCUSSION

In this paper, we studied an explicit QTL model using a forward simulation framework to assess the power of short-term E&R experiments in detecting QTLs. We found that the expected power of such experiments depends strongly on the genetic architecture underlying the trait of interest. This relationship is complex and often unintuitive, and we demonstrated that few results hold universally, except perhaps for the fact that as the number of QTLs affecting a trait increases, the power to detect them always decreases (Figure 3). Other results are often conditional on the false positive rate, the precise way in which power is measured, and various aspects of trait architecture. For example, we found that the clustering of QTLs along a chromosome decreases detection power when there are fewer QTLs, but increases power at low false positive rates when there are more QTLs (Figure 4). An exponential distribution in effect size lowers the proportion of QTLs detected compared to when effect sizes are equal at all QTLs, but increases power when QTLs are weighted by their effect sizes (i.e. we have a better chance at detecting a substantial portion of the genetic basis of a trait, Figure 5). When minor alleles at QTLs are skewed towards lower frequencies, detection power is generally decreased, but when they start at higher frequencies, detection power is increased only when it is measured by the proportion of QTLs detected (Figure 6). When derived alleles are dominant, detection power is increased at higher false positive rates but decreased at lower (Figure 7). When they are recessive, detection power is decreased only when it is evaluated by the proportion of QTLs detected (Figure 7). Epistasis tends to always decrease power, although a lot of variation exists among different types of epistasis (Figure 8). In spite of all these complexities, we found that when the trait of interest is controlled primarily by a few unlinked QTLs that show little dominance or epistatic effects and start at relatively high frequency, E&R experiments spanning as few as 4 generations with a population as small as 1000 individuals can still be a viable tool for QTL detection (Figure 3 & S4).

From the perspective of researchers who plan to perform E&R studies, a key question might be which experimental designs could optimize detection power (Kofler and Schlötterer 2014; Baldwin-Brown *et al.* 2014; Kessner and Novembre 2015). Our results suggest that there is not a single optimal strategy that is always expected to work best for all trait architectures. Instead, what the best strategy is can depend critically on the genetic architecture of the trait. For example, in our standard model with 10 QTLs, detection power can be significantly improved if the selection experiment were to be run for an additional 5 generations, since many QTLs are still segregating at intermediate frequency after just 4 generations of selection. However, the gain in power will be minimal if derived alleles are either dominant or recessive, since during the additional generations, selection would become less effective in the dominant case, while most of the genetic variance will already have been depleted in the recessive case (Figure 9). When researchers have some information on the expected trait architecture, different experimental designs can be simulated to find the optimal setup for the given architecture. Without such *a priori* information, they may first need to simulate a range of architectures to obtain a general sense of the range of power that could be achieved and what types of architectures are likely to be detectable under different experimental designs.

**Figure 9.**
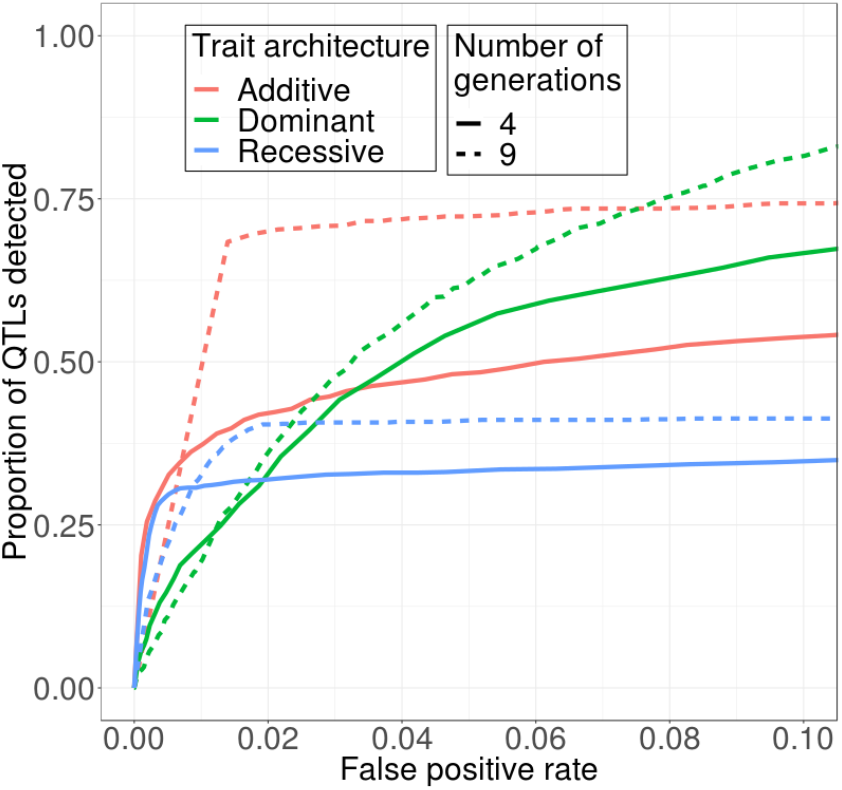
The choice of optimal experimental design depends on the underlying trait architecture. Extending the selection experiment from 4 to 9 generations substantially increases detection power when the derived allele is codominant, but such effect is minimal when the derived allele is either recessive or dominant. Models with 10 QTLs are used. Additive: heterozygotes express intermediate phenotype. Dominant: heterozygotes express the same phenotype as derived homozygotes. Recessive: heterozygotes express the same phenotype as ancestral homozygotes.

Our results highlight the complex nature of the mechanisms involved, and thus the crucial role simulations can play in assessing the expected detection power and optimizing the experimental design. To this end, we provide a flexible simulation framework that is highly customizable with regard to trait architecture, experimental design, and parameters of the genomic background (described in the Supplementary Materials). While several specialized tools have already been designed for such applications (Neuenschwander *et al.* 2008; Zanini and Neher 2012; Kofler and Schlötterer 2014; Kessner and Novembre 2015; Vlachos and Kofler 2018), our approach provides key advantages by implementing its simulations in the flexible SLiM framework (Haller and Messer 2016, 2019). SLiM constitutes one of the most widely used and well-tested frameworks for forward genetic simulation to date. This popularity is grounded in several attributes. First, simulations are fully scriptable, allowing the user to model a wide range of evolutionary scenarios that can include high levels of genetic and ecological realism. Second, the comprehensive graphical user interface enables interactive model development and allows for easy visual debugging and hands-on model exploration. Third, the underlying simulation engine has been highly optimized over the years, and these speed benefits are now inherited for free by any new simulation model built in the SLiM framework. For example, with the recently added capability for tree sequence recording (Haller *et al.* 2019), forward simulations on a genome-wide scale can now be conducted in runtimes that are on par with efficient coalescent-based simulations, enabling applications such as approximate Bayesian computation (ABC) for parameter inference (Csilléry *et al.* 2010). All SLiM configuration scripts developed in this study, together with a comprehensive documentation explaining how these scripts may be adjusted for custom scenarios, are provided in the Supplementary Materials.

One perhaps surprising result of our analyses was that a simple summary statistic, *D*, had comparable and often better performance than more sophisticated, model-based methods explicitly devised for detecting positive selection from time-series data. This is presumably because these methods were built on the assumption that selection produces independent selective sweeps at individual QTLs. However, when selection operates on a polygenic trait, allele frequency trajectories at its QTLs will often be quite distinct from those expected in a model of individual sweeps with fixed selection coefficients (Burke *et al.* 2010; Kessner and Novembre 2015; Franssen *et al.* 2017). In addition, QTLs are usually not freely recombining, further complicating allele frequency trajectories due to interference effects. Simple summary statistics that do not rely on detailed, yet potentially inaccurate, assumptions about the precise form of the temporal allele frequency trajectories may actually work better in these scenarios than the more sophisticated model-based methods. However, we recognize that when only a single selection line is created and the trait under selection has a simple genetic architecture, model-based methods may still be able to outperform *D*, especially when their parameters are optimally tuned and they can take advantage of their ability to analyze the full allele frequency trajectories, instead of just the absolute change in allele frequencies over the course of the experiment as measured by the *D* statistic (Figure S3).

Compared with earlier power analyses of E&R experiments, detection power was generally lower in our selection model. This can be attributed to the following factors. First, selection is quite extreme in our model, while the length of the experiment is very short. Such an experimental design would likely be considered as ineffective when working with an insect species such as flies, but may be the only realistic choice for larger organisms with longer generation times. Second, we assumed a higher level of linkage disequilibrium and thus stronger interference effects than previous studies that focused on insect systems. We also note that for simplicity we set heritability to a value of one in our model, which should generally lead to an overestimation of power (Kessner and Novembre 2015). This can easily be modified in our SLiM simulations if a more realistic estimation of power for a specific organism and trait of interest is desired.

## CONCLUSION

Overall, we find that short E&R experiments with strong truncating selection can provide some utility in identifying the genetic basis of the selected trait, especially if the goal is to detect those QTLs that contribute most to the observed trait variance in the population (i.e. large effect QTLs present at high population frequency). However, we have also shown that detection power can vary substantially with the genetic architecture of the trait. This presents a problem of circular reasoning, because the architecture of the trait will likely be unknown prior to the experiment yet the power to identify its genetic basis should be biased against certain classes of QTLs (e.g. those that are dominant/recessive, have epistatic interactions, or are present at low initial frequencies). Thus, some caution is warranted when trying to make general conclusions about the architecture of the selected trait, based solely on the subset of QTLs that were identified in an E&R experiment. Future studies will hopefully improve our understanding of what types of trait architectures are more prevalent in nature and thereby help us build better priors for the interpretation of E&R experiments.

## ACKNOWLEDGMENTS

We would like to thank John Novembre, Kevin Thornton, and Matt Hare, for helpful comments and discussion. This work was supported by Cornell University (a Presidential Life Science Fellowship to RNL and a Center for Vertebrate Genomics Seed Grant to NOT and PWM), and the National Science Foundation (NSF OCE-1756316 to NOT).

## REFERENCES

Backström N., W. Forstmeier, H. Schielzeth, H. Mellenius, K. Nam, et al., 2010 The recombination landscape of the zebra finch *Taeniopygia guttata* genome. Genome Res. 20: 485–495. https://doi.org/10.1101/gr.101410.109

Baldwin-Brown J. G., A. D. Long, and K. R. Thornton, 2014 The power to detect quantitative trait loci using resequenced, experimentally evolved populations of diploid, sexual organisms. Mol. Biol. Evol. 31: 1040–1055. https://doi.org/10.1093/molbev/msu048

Barrett R. D. H., A. Paccard, T. M. Healy, S. Bergek, P. M. Schulte, et al., 2011 Rapid evolution of cold tolerance in stickleback. Proc. R. Soc. B Biol. Sci. 278: 233–238. https://doi.org/10.1098/rspb.2010.0923

Barrett R. D. H., S. Laurent, R. Mallarino, S. P. Pfeifer, C. C. Y. Xu, et al., 2019 Linking a mutation to survival in wild mice. Science 363: 499–504. https://doi.org/10.1126/science.aav3824

Barrick J. E., D. S. Yu, S. H. Yoon, H. Jeong, T. K. Oh, et al., 2009 Genome evolution and adaptation in a long-term experiment with *Escherichia coli*. Nature 461: 1243–1247. https://doi.org/10.1038/nature08480

Barton N. H., and M. Turelli, 1989 Evolutionary quantitative genetics: How little do we know? Annu. Rev. Genet. 23: 337–370. https://doi.org/10.1146/annurev.ge.23.120189.002005

Burger J. C., M. A. Chapman, and J. M. Burke, 2008 Molecular insights into the evolution of crop plants. Am. J. Bot. 95: 113–122. https://doi.org/10.3732/ajb.95.2.113

Burke M. K., J. P. Dunham, P. Shahrestani, K. R. Thornton, M. R. Rose, et al., 2010 Genome-wide analysis of a long-term evolution experiment with *Drosophila*. Nature 467: 587–590. https://doi.org/10.1038/nature09352

Cai H., and H. Morishima, 2002 QTL clusters reflect character associations in wild and cultivated rice. Theor. Appl. Genet. 104: 1217–1228. https://doi.org/10.1007/s00122-001-0819-7

Castro J. P. L., M. N. Yancoskie, M. Marchini, S. Belohlavy, M. Kučka, et al., 2018 An integrative genomic analysis of the Longshanks selection experiment for longer limbs in mice. bioRxiv 378711. https://doi.org/10.1101/378711

Chan Y. F., F. C. Jones, E. McConnell, J. Bryk, L. Bünger, et al., 2012 Parallel selection mapping using artificially selected mice reveals body weight control loci. Curr. Biol. 22: 794–800. https://doi.org/10.1016/j.cub.2012.03.011

Charlesworth D., and B. Charlesworth, 1979 Selection on recombination in clines. Genetics 91: 581–589.

Chen X., R. Kuja-Halkola, I. Rahman, J. Arpegård, A. Viktorin, et al., 2015 Dominant genetic variation and missing heritability for human complex traits: Insights from twin versus genome-wide common SNP models. Am. J. Hum. Genet. 97: 708–714. https://doi.org/10.1016/j.ajhg.2015.10.004

Chevalet C., 1994 An approximate theory of selection assuming a finite number of quantitative trait loci. Genet. Sel. Evol. GSE 26: 379–400. https://doi.org/10.1186/1297-9686-26-5-379

Christie M. R., M. L. Marine, S. E. Fox, R. A. French, and M. S. Blouin, 2016 A single generation of domestication heritably alters the expression of hundreds of genes. Nat. Commun. 7: 10676. https://doi.org/10.1038/ncomms10676

Coltman D. W., P. O’Donoghue, J. T. Jorgenson, J. T. Hogg, C. Strobeck, et al., 2003 Undesirable evolutionary consequences of trophy hunting. Nature 426: 655–658. https://doi.org/10.1038/nature02177

Conover D. O., and S. B. Munch, 2002 Sustaining fisheries yields over evolutionary time scales. Science 297: 94–96. https://doi.org/10.1126/science.1074085

Csilléry K., M. G. B. Blum, O. E. Gaggiotti, and O. François, 2010 Approximate Bayesian Computation (ABC) in practice. Trends Ecol. Evol. 25: 410–418. https://doi.org/10.1016/j.tree.2010.04.001

Dowle M., A. Srinivasan, J. Gorecki, M. Chirico, P. Stetsenko, et al., 2019 data.table: Extension of “data.frame.”

Fallahsharoudi A., N. de Kock, M. Johnsson, L. Bektic, S. J. K. A. Ubhayasekera, et al., 2017 Genetic and targeted eQTL mapping reveals strong candidate genes modulating the stress response during chicken domestication. G3 Genes Genomes Genet. 7: 497–504. https://doi.org/10.1534/g3.116.037721

Ferrer-Admetlla A., C. Leuenberger, J. D. Jensen, and D. Wegmann, 2016 An approximate Markov model for the Wright-Fisher diffusion and its application to time series data. Genetics 203: 831–846. https://doi.org/10.1534/genetics.115.184598

Foll M., H. Shim, and J. D. Jensen, 2015 WFABC: a Wright–Fisher ABC-based approach for inferring effective population sizes and selection coefficients from time-sampled data. Mol. Ecol. Resour. 15: 87–98. https://doi.org/10.1111/1755-0998.12280

Franssen S. U., R. Kofler, and C. Schlötterer, 2017 Uncovering the genetic signature of quantitative trait evolution with replicated time series data. Heredity 118: 42–51. https://doi.org/10.1038/hdy.2016.98

Fuller R. C., C. F. Baer, and J. Travis, 2005 How and When Selection Experiments Might Actually be Useful. Integr. Comp. Biol. 45: 391–404. https://doi.org/10.1093/icb/45.3.391

Garland T., and M. R. Rose, 2009 Experimental evolution: concepts, methods, and applications of selection experiments. University of California Press.

Gibson G., 2012 Rare and common variants: twenty arguments. Nat. Rev. Genet. 13: 135–145. https://doi.org/10.1038/nrg3118

Gutierrez A. P., J. M. Yáñez, and W. S. Davidson, 2016 Evidence of recent signatures of selection during domestication in an Atlantic salmon population. Mar. Genomics 26: 41–50. https://doi.org/10.1016/j.margen.2015.12.007

Haller B. C., and P. W. Messer, 2016 SLiM 2: Flexible, interactive forward genetic simulations. Mol. Biol. Evol. msw211. https://doi.org/10.1093/molbev/msw211

Haller B. C., J. Galloway, J. Kelleher, P. W. Messer, and P. L. Ralph, 2019 Tree-sequence recording in SLiM opens new horizons for forward-time simulation of whole genomes. Mol. Ecol. Resour. 19: 552–566. https://doi.org/10.1111/1755-0998.12968

Haller B. C., and P. W. Messer, 2019 SLiM 3: Forward genetic simulations beyond the Wright-Fisher model. Mol. Biol. Evol. 36: 632–637. https://doi.org/10.1093/molbev/msy228

Hansen T. F., 2006 The evolution of genetic architecture. Annu. Rev. Ecol. Evol. Syst. 37: 123–157. https://doi.org/10.1146/annurev.ecolsys.37.091305.110224

Hill W. G., and A. Robertson, 1966 The effect of linkage on limits to artificial selection. Genet. Res. 8: 269–294. https://doi.org/10.1017/S0016672300010156

Hill W. G., and A. Caballero, 1992 Artificial selection experiments. Annu. Rev. Ecol. Syst. 23: 287–310. https://doi.org/10.1146/annurev.es.23.110192.001443

Houde A. E., 1994 Effect of artificial selection on male colour patterns on mating preference of female guppies. Proc R Soc Lond B 256: 125–130. https://doi.org/10.1098/rspb.1994.0059

Johansson A. M., M. E. Pettersson, P. B. Siegel, and Ö. Carlborg, 2010 Genome-wide effects of long-term divergent selection. PLOS Genet. 6: e1001188. https://doi.org/10.1371/journal.pgen.1001188

Kelly J. K., and K. A. Hughes, 2019 Pervasive linked selection and intermediate-frequency alleles are implicated in an evolve-and-resequencing experiment of *Drosophila simulans*. Genetics 211: 943–961. https://doi.org/10.1534/genetics.118.301824

Kessner D., and J. Novembre, 2015 Power analysis of artificial selection experiments using efficient whole genome simulation of quantitative traits. Genetics 199: 991–1005. https://doi.org/10.1534/genetics.115.175075

Kirkpatrick M., 2006 Chromosome inversions, local adaptation and speciation. Genetics 173: 419–434. https://doi.org/10.1534/genetics.105.047985

Kofler R., and C. Schlötterer, 2014 A guide for the design of evolve and resequencing studies. Mol. Biol. Evol. 31: 474–483. https://doi.org/10.1093/molbev/mst221

Lang G. I., D. P. Rice, M. J. Hickman, E. Sodergren, G. M. Weinstock, et al., 2013 Pervasive genetic hitchhiking and clonal interference in forty evolving yeast populations. Nature 500: 571–574. https://doi.org/10.1038/nature12344

Long A., G. Liti, A. Luptak, and O. Tenaillon, 2015 Elucidating the molecular architecture of adaptation via evolve and resequence experiments. Nat. Rev. Genet. 16: nrg3937. https://doi.org/10.1038/nrg3937

Mackay T. F., 2009 Q&A: Genetic analysis of quantitative traits. J. Biol. 8: 23. https://doi.org/10.1186/jbiol133

Mackay T. F. C., E. A. Stone, and J. F. Ayroles, 2009 The genetics of quantitative traits: challenges and prospects. Nat. Rev. Genet. 10: 565–577. https://doi.org/10.1038/nrg2612

Mackay T. F. C., 2014 Epistasis and quantitative traits: using model organisms to study gene-gene interactions. Nat. Rev. Genet. 15: 22–33. https://doi.org/10.1038/nrg3627

Malaspinas A.-S., 2016 Methods to characterize selective sweeps using time serial samples: an ancient DNA perspective. Mol. Ecol. 25: 24–41. https://doi.org/10.1111/mec.13492

Neuenschwander S., F. Hospital, F. Guillaume, and J. Goudet, 2008 quantiNemo: an individual-based program to simulate quantitative traits with explicit genetic architecture in a dynamic metapopulation. Bioinformatics 24: 1552–1553. https://doi.org/10.1093/bioinformatics/btn219

Oortmerssen G. A. van, and T. C. M. Bakker, 1981 Artificial selection for short and long attack latencies in wild *Mus musculus domesticus*. Behav. Genet. 11: 115–126. https://doi.org/10.1007/BF01065622

Orr H. A., 1998 The population genetics of adaptation: The distribution of factors fixed during adaptive evolution. Evolution 52: 935–949. https://doi.org/10.1111/j.1558-5646.1998.tb01823.x

Otto S. P., and C. D. Jones, 2000 Detecting the undetected: estimating the total number of loci underlying a quantitative trait. Genetics 156: 2093–2107.

Parts L., F. Cubillos, J. Warringer, K. Jain, F. Salinas, et al., 2011 Revealing the genetic structure of a trait by sequencing a population under selection. Genome Res. gr.116731.110. https://doi.org/10.1101/gr.116731.110

Pigeon G., M. Festa-Bianchet, D. W. Coltman, and F. Pelletier, 2016 Intense selective hunting leads to artificial evolution in horn size. Evol. Appl. 9: 521–530. https://doi.org/10.1111/eva.12358

Roesti M., D. Moser, and D. Berner, 2013 Recombination in the threespine stickleback genome—patterns and consequences. Mol. Ecol. 22: 3014–3027. https://doi.org/10.1111/mec.12322

Rubin C.-J., M. C. Zody, J. Eriksson, J. R. S. Meadows, E. Sherwood, et al., 2010 Whole-genome resequencing reveals loci under selection during chicken domestication. Nature 464: 587–591. https://doi.org/10.1038/nature08832

Sadowska E. T., K. Baliga-Klimczyk, K. M. Chrząścik, and P. Koteja, 2008 Laboratory model of adaptive radiation: A selection experiment in the bank vole. Physiol. Biochem. Zool. 81: 627–640. https://doi.org/10.1086/590164

Schlötterer C., R. Kofler, E. Versace, R. Tobler, and S. U. Franssen, 2015 Combining experimental evolution with next-generation sequencing: a powerful tool to study adaptation from standing genetic variation. Heredity 114: 431–440. https://doi.org/10.1038/hdy.2014.86

Seabra S. G., I. Fragata, M. A. Antunes, G. S. Faria, M. A. Santos, et al., 2018 Different genomic changes underlie adaptive evolution in populations of contrasting history. Mol. Biol. Evol. 35: 549–563. https://doi.org/10.1093/molbev/msx247

Shao H., L. C. Burrage, D. S. Sinasac, A. E. Hill, S. R. Ernest, et al., 2008 Genetic architecture of complex traits: Large phenotypic effects and pervasive epistasis. Proc. Natl. Acad. Sci. 105: 19910–19914. https://doi.org/10.1073/pnas.0810388105

Shifman S., J. T. Bell, R. R. Copley, M. S. Taylor, R. W. Williams, et al., 2006 A high-resolution single nucleotide polymorphism genetic map of the mouse genome. PLOS Biol. 4: e395. https://doi.org/10.1371/journal.pbio.0040395

Slatkin M., 1975 Gene glow and selection in a two-locus system. Genetics 81: 787–802.

Smith J. M., and J. Haigh, 1974 The hitch-hiking effect of a favourable gene. Genet. Res. 23: 23–35. https://doi.org/10.1017/S0016672300014634

Stetter M. G., K. Thornton, and J. Ross-Ibarra, 2018 Genetic architecture and selective sweeps after polygenic adaptation to distant trait optima. PLOS Genet. 14: e1007794. https://doi.org/10.1371/journal.pgen.1007794

Swain D. P., A. F. Sinclair, and J. M. Hanson, 2007 Evolutionary response to size-selective mortality in an exploited fish population. Proc. R. Soc. Lond. B Biol. Sci. 274: 1015–1022. https://doi.org/10.1098/rspb.2006.0275

Tenaillon O., A. Rodríguez-Verdugo, R. L. Gaut, P. McDonald, A. F. Bennett, et al., 2012 The molecular diversity of adaptive convergence. Science 335: 457–461. https://doi.org/10.1126/science.1212986

Teotónio H., S. Estes, P. C. Phillips, and C. F. Baer, 2017 Experimental evolution with *Caenorhabditis* nematodes. Genetics 206: 691. https://doi.org/10.1534/genetics.115.186288

Therkildsen N. O., J. Hemmer-Hansen, T. D. Als, D. P. Swain, M. J. Morgan, et al., 2013 Microevolution in time and space: SNP analysis of historical DNA reveals dynamic signatures of selection in Atlantic cod. Mol. Ecol. 22: 2424–2440. https://doi.org/10.1111/mec.12260

Turner T. L., A. D. Stewart, A. T. Fields, W. R. Rice, and A. M. Tarone, 2011 Population-based resequencing of experimentally evolved populations reveals the genetic basis of body size variation in *Drosophila melanogaster*. PLOS Genet. 7: e1001336. https://doi.org/10.1371/journal.pgen.1001336

Uusi-Heikkilä S., T. Sävilammi, E. Leder, R. Arlinghaus, and C. R. Primmer, 2017 Rapid, broad-scale gene expression evolution in experimentally harvested fish populations. Mol. Ecol. 26: 3954–3967. https://doi.org/10.1111/mec.14179

Vlachos C., and R. Kofler, 2018 MimicrEE2: Genome-wide forward simulations of Evolve and Resequencing studies. PLOS Comput. Biol. 14: e1006413. https://doi.org/10.1371/journal.pcbi.1006413

Wickham H., and RStudio, 2017 tidyverse: Easily install and load the “tidyverse.”

Wilke C. O., and RStudio, 2019 cowplot: Streamlined plot theme and plot annotations for “ggplot2.”

Wong A. K., A. L. Ruhe, B. L. Dumont, K. R. Robertson, G. Guerrero, et al., 2010 A comprehensive linkage map of the dog genome. Genetics 184: 595–605. https://doi.org/10.1534/genetics.109.106831

Zanini F., and R. A. Neher, 2012 FFPopSim: an efficient forward simulation package for the evolution of large populations. Bioinformatics 28: 3332–3333. https://doi.org/10.1093/bioinformatics/bts633

Zhou D., N. Udpa, M. Gersten, D. W. Visk, A. Bashir, et al., 2011 Experimental selection of hypoxia-tolerant *Drosophila melanogaster*. Proc. Natl. Acad. Sci. 201010643. https://doi.org/10.1073/pnas.1010643108

