## Supplementary Materials for "The Effects of Quantitative Trait Architecture on Detection Power in Artificial Selection Experiments"

### Simulation pipeline instructions

Here we provide a customizable tool for simulation of evolve and resequence experiments on a quantitative trait to evaluate their power for QTL detection. The simulation is carried out in the program SLiM and the data analysis is conducted in R.

Below is our suggested workflow. Each step in the workflow has to be completed before the next can start.

For the simulation of E&R experiment:

1. Create neutral populations representing the population prior to the selection experiment. To do this, read and edit ShellScripts/Burnin.sh. Then, run Burnin.sh.
2. Establish the trait architecture and simulate the selection experiment. For this, read and edit ShellScripts/Selection.sh, then run the script. This step can be repeated to simulate different combinations of trait architecture and experimental design. These can be created using the same burn-in.

For the power analysis in QTL detection:

3. Compile the SLiM outputs from each simulation in step 2, and (optionally) create input files for WFABC and ApproxWF. To do this, read, edit, and run RScripts/Compile.R
4. Calculate the power and false positive rate for each simulation. For this, read, edit, and run RScripts/Analysis.R
5. Plot ROC curve comparisons by combining the ROC tables from simulations of different trait architecture or experimental design given by step 4. You can use RScripts/PlotROC.R to do this. Alternatively, you can also create your own R scripts, since this step highly depends on what your specific comparison is.

Note:

To adjust most of the variables, you will not need to edit the SLiM script and can directly input them through Burnin.sh and Selection.sh. Below is a list of these variables.

| **Trait architecture** | **Population parameters** | **Experimental design setting** |
|---|---|---|
| Number of QTLs | Population size* | Sample size |
| Position of QTLs | Number of chromosomes | Length of experiment* |
| Effect sizes of QTLs | Length of chromosomes | Mode of selection |
| Starting frequency of QTLs | Recombination rate | Strength and direction of selection |
| Dominance | Nucleotide diversity | |
| Pairwise epistasis | | |

* To modify these two variables, one single value will need to be changed in the .slim files. See the scripts for details.

There are some variables and scenarios that are not explicitly incorporated in the SLiM script but that can be conveniently implemented by editing SlimScripts/Burnin.slim and SlimScripts /Selection.slim. These variables and scenarios include:

- heritability values other than 1 (see the SLiM manual Chapter 13.4)
- selection modes other than truncating selection (see the SliM manual Chapter 13.1), pleiotropy (see the SLiM manual Chapter 13.5)
- population structure (see the SLiM manual Chapter 5.2)
- demographic history prior to the selection experiment (see the SLiM manual Chapter 5)
- known genotypic data of the experimental population (see the SLiM manual Chapter 18.12)

For data analyses and visualizations (step 3-5), it might be more efficient if you write your own scripts from scratch.

If WFABC and/or ApproxWF are used, they should to be run between step 3 and 4.

Please post an issue on GitHub or contact us at with any problems or questions.

### Simulation pipeline scripts

*Burn-in*

#### Burnin.sh:

```
## This script is used to generate neutral populations used for the selection experiment.
## IMPORTANT: Change the output directory in the first step and the variables in the second step
before running.
## Run the script on server using: nohup bash Burnin.sh > Burnin.nohup &
## Note: the number of generations to be included in the burn-in process depends on the
population size (typically ten times the population size) and needs to be changed in the
Burnin.slim file. Please refer to the notes in that file.

## Step 1: Create directories to store the outputs. Make sure to change the directory name in the
first line.
cd /fs/cbsubscb10/storage/rl683/TemporalScan/Simulations # Change this to a directory where you
want to store all you simulation outputs.
mkdir Burnin
cd Burnin
for k in {1..100} # Number of simulation replicates that you want to create.
do
    mkdir 'SimRep'$k
done
cd ..

## Step 2: Run burnin using SLiM 2. Variables inside the loop are all customizable and can be
changed as desired.
for k in {1..100} # Set the number of simulation replicates that you want to create.
do
    ~/Program/SLiM/bin/slim \ # slim directory
    -d SimRepID=$k  \  # SimuRepID = Simulation Replicate ID
    -d Mu=2e-8 \ # Mu = mutation rate
    -d RecRate=1e-8 \ # RecRate = recombination rate (change the slim script if simulating
multiple chromosomes)
    -d LCh=30000000 \ # LCh = length of chromosome
    -d BurninSize=1000 \ # BurninSize = size of the burnin populations (IF THIS NEEDS TO BE
CHANGED, MAKE SURE TO CHANGE MUTATION RATE, RECOMBINATION RATE, AND NUMBER OF GENERATIONS
ACCORDINGLY)
    -d "BurninPath='/fs/cbsubscb10/storage/rl683/TemporalScan/Simulations/Burnin/'" \ #
BurninPath = path to the burnin files
    -d "BurninFilename='Burnin.txt'" \ # BurninFilename = name of the burnin file
    /fs/cbsubscb10/storage/rl683/TemporalScan/SlimScripts/Burnin.slim # Directory to the
Burnin.slim file included in the simulation tool.
done
```

#### Burnin.slim:

```
// The number of generations in the burn-in process depends on the population size (typically ten
times the population size). If it needs to be changed, change the "10000" value in beginning of
the third and fourth section to the number of generations desired.

initialize() {
        // set mutation rate
        initializeMutationRate(Mu);

        // neutral mutations
        initializeMutationType("m1", 0.5, "f", 0.0);
        initializeGenomicElementType("g1", m1, 1.0);
```

```
        // set up the chromosome
        initializeGenomicElement(g1, 0, LCh-1);

        // set recombination rate
        initializeRecombinationRate(RecRate, LCh-1);
}

1 early() {
        // define a population
        sim.addSubpop("p1", BurninSize);
}

10000 late(){
        // save the output of the burn-in
        sim.outputFull(BurninPath + "SimRep" + SimRepID + "/" + BurninFilename);
}

s1 10000 late() {
}
```

### *Selection experiment*

#### Selection.sh:

```
## This script is used to establish quantitative trait architectures and perform artificial selection experiments on the populations generated using the Burnin.sh script.
## IMPORTANT: Change the output directory in the first step and the variables in the second step before running.
## Run the script on a server using: nohup bash Selection.sh > Selection.nohup &

## Step 1: Create directories to store the outputs. Make sure to change the directory names in the first three lines.
cd /fs/cbsubscb10/storage/rl683/TemporalScan/Simulations/ # Change this to a directory where you want to store all your simulation outputs.
mkdir NQTL10 # Change this to what you want to name this particular quantitative trait architectures and/or the experimental design that you are simulating.
cd NQTL10 # Same as above
for k in {1..100} # Number of simulation replicates that you want to create.
do
    mkdir 'SimRep'$k
    cd 'SimRep'$k
    for j in {1..10}
    do
        mkdir 'ExpRepPlus'$j
        mkdir 'ExpRepMinus'$j
    done
    cd ..
done
cd ..

## Step 2: Run the selection experiment using SLiM 2. Variables inside the loop are all customizable and can be changed as desired.
## It is recommended if you copy this shell script for each of the trait architecture x experimental combinations that you want to test and make changes in the new script.
## For example, when simulating the scenario with 100 QTLs, copy this file and rename it as NQTL100.sh. Set the NQTL variable to be 100 and run it with "nohup bash NQTL100.sh > NQTL100.nohup &"
## Note: The number of generations in the selection experiment can only be changed in the Selection.slim script.
```

```
## Note: Depending on the population size, more or fewer burn-in generations might have been needed. If that is the case, edit the Selection.slim file as instructed in the file.

for k in {1..100} # Set the number of simulation replicates that you want to create.
do
    echo $k
    for j in {1..1} # Set the number of experimental replications.
    do
        echo $j
        for i in {T,F} # Set the direction of selection (F if selecting the larger phenotype, T otherwise, T,F is both directions are selected)
        do
            echo $i
            ~/Program/SLiM/bin/slim \ # slim directory
            -d SimRepID=$k  \ # SimuRepID = Simulation Replicate ID
            -d ExpRepID=$j \ # ExpRepID = Experiment replicate ID
            -d Direction=$i \ # Direction = direction of selection; set this at the line "for i in {T,F}" above
            -d "BurninPath='/fs/cbsubscb10/storage/rl683/TemporalScan/Simulations/Burnin/'" \ # BurninPath = path to the burnin files
            -d "BurninFilename='Burnin.txt'" \ # BurninFilename = name of the burnin file
            -d LCh=30000000 \ # LCh = length of chromosome (CHANGE THIS ONLY WHEN THE BURNIN USES A DIFFERENT CHROMOSOME SIZE)
            -d RecRate=1e-8 \ # RecRate = recombination rate (change the slim script if simulating multiple chromosomes)
            -d SampleSize=50 \ # SampleSize = number of individuals to sample each generation
            -d NQTL=10 \ # NQTL = number of QTLs, (even number is recommended when the number is small)
            -d ESMean=1.0 \ # ESMean = absolute value of mean effect size
            -d "ESDist='f'" \ # ESDist = effect size distribution("f" for fixed or "e" for exponential),
            -d LowFreq=F \ # LowFreq = starting frequency preference (T if selecting for lower frequency, F if selecting for higher frequency or random)
            -d FreqBound=0.0 \ # FreqBound = frequency bound (0.0~0.5, 0.0 if starting frequency is random)
            -d LowerPosBound=0 \ # LowerPosBound = lower position bound (0 if random)
            -d UpperPosBound=29999999 \ # UpperPosBound = upper position bound (LCh-1 if random)
            -d D=0.5 \ # D = dominance coefficient (0.0~1.0, 1.0 for mutant being completely dominant and 0.0 for wildtype to being completely dominant.)
            -d Epistasis=F \ # Epistasis = F if there is no epistasis, T otherwise
            -d "EpiSce=c(0,1,2,1,2,3,2,3,4)" \ # EpiSce = epistasis scenario; vector of size 9; first element must be 0; needs to be defined if Epistasis ==T; the nine value corresponds to phenotypes of genotypes in the following order: c(aabb, Aabb, AAbb, aaBb, AaBb, AABb, aaBB, AaBB, AABB); e.g. c(0,1,2,1,2,3,2,3,4) when there is no epistasis.
            -d PopSize=1000 \ # PopSize = population size (can only downsample)
            -d SelectedSize=100 \ # SelectedSize = number of selected individuals in each generation
            -d "OutPath='/fs/cbsubscb10/storage/rl683/TemporalScan/Simulations/NQTL10/'" \ # OutPath = output path
            /fs/cbsubscb10/storage/rl683/TemporalScan/SlimScripts/Selection.slim # Directory to the Selection.slim file included in the simulation tool.
        done
    done
done
```

### Selection.slim:

```
// The first variable that might need to be edited in this slim file is number of generations in the selection experiment. It is defined in the last section of this file, in the line "10000:10010 late() {". In this example, the selection experiment lasts 11 generations. If you want to set it to 100 generation, for example, this line should become "10000:10099 late() {"
```

```
// Depending on the population size, more or fewer burn-in generations might have been needed.
Therefore, the second variable that might need to be modified is the number 10000 in the
beginning of each section; this number should be changed to reflect the number of generations in
the burn-in.

initialize() {
        initializeMutationRate(0);

        // neutral mutations
        initializeMutationType("m1", 0.5, "f", 0.0);
        initializeGenomicElementType("g1", m1, 1.0);
        m1.convertToSubstitution = F;

        // mutations representing alleles in QTLs
        initializeMutationType("m2", D, "f", 0.0);
        m2.convertToSubstitution = F;

        // a mutation type used to evaluate phenotypic fitness
        initializeMutationType("m3", 0.5, "f", 0.0);
        m3.convertToSubstitution = F;

        // set up the chromosome
        initializeGenomicElement(g1, 0, LCh-1);

        // set recombination rate; change this if several chromosomes are simulated
        initializeRecombinationRate(RecRate, LCh-1);

}

10000 late() {
        // read the burn-in
        sim.readFromPopulationFile(BurninPath + "SimRep" + SimRepID + "/" + BurninFilename);

        // give an m3 mutation to every individual, for phenotypic fitness
        sim.subpopulations.genomes.addNewMutation(m3, 0.0, 0);

        // sanity check
        // print(size(unique(sim.subpopulations.genomes.mutationsOfType(m1))));
        // print(size(unique(sim.subpopulations.genomes.mutationsOfType(m1).position)));

        // get all SNPs within an starting frequency interval
        if (LowFreq == T)

        FreqencyConstrainedMutations=unique(sim.subpopulations.genomes.mutationsOfType(m1))[
                which(sim.mutationFrequencies(p1,
unique(sim.subpopulations.genomes.mutationsOfType(m1))) <= FreqBound |
                sim.mutationFrequencies(p1,
unique(sim.subpopulations.genomes.mutationsOfType(m1))) >= (1-FreqBound))];
        else

        FreqencyConstrainedMutations=unique(sim.subpopulations.genomes.mutationsOfType(m1))[
                which(sim.mutationFrequencies(p1,
unique(sim.subpopulations.genomes.mutationsOfType(m1))) >= FreqBound &
                sim.mutationFrequencies(p1,
unique(sim.subpopulations.genomes.mutationsOfType(m1))) <= (1-FreqBound))];

        // get all SNPs within one region of a chromosome
        ClusteredMutations=unique(FreqencyConstrainedMutations)
        [which(FreqencyConstrainedMutations.position >= LowerPosBound &
        FreqencyConstrainedMutations.position <= UpperPosBound)];

        // set seed so that QTLs remain the same within one simulation replication
        setSeed(SimRepID);

        // sample QTLs at random after the frequency and position constraints are imposed
        QTL=sample(ClusteredMutations,NQTL);
        QTL.setMutationType(m2);

        // assign positive effect sizes to all QTLs first
```

```
        if (ESDist == "f")
                for (IndividualQTL in QTL)
                {
                IndividualQTL.setSelectionCoeff(ESMean);
                }
        else if (ESDist == "e")
                for (IndividualQTL in QTL)
                {
                IndividualQTL.setSelectionCoeff(rexp(1,ESMean));
                }

        // assign negative effect sizes to half of the QTLs
        Negative = sample(0:(NQTL-1), asInteger(NQTL/2));
        print(Negative);
        NegativeQTL = QTL[Negative];
        for (IndividualQTL in NegativeQTL)
                {
                IndividualQTL.setSelectionCoeff(-(IndividualQTL.selectionCoeff));
                }
        // if there is epistasis, randomly set up epistatic pairs
        if (Epistasis == T)
                QTL.tag = 1:NQTL;

        // set experimental population size (only takes effect in the next generation)
    p1.setSubpopulationSize(PopSize);

    // more sanity checks
        // print(sim.subpopulations);
        // print(size(sim.subpopulations.individuals));

        // set different seeds for different experimental replications for later steps
    setSeed(SimRepID*ExpRepID);

        // even more checks
        // print(QTL);
        // print(QTL.position);
        // print(sim.mutationFrequencies(p1, QTL));
}

10000: late() {
        // define generation number
        GenID = sim.generation-9999;

        // define individual tags as phenotypes for the additive effects of QTLs while accounting 
for dominance
        for (individual in sim.subpopulations.individuals)
        {
                        if (Epistasis == F)
                        {
                                // extract sites when the mutant alleles are hehomozygous or 
heterozygous as needed for dominance scenarios
                                Homo = 
individual.genomes[0].mutationsOfType(m2)[match(individual.genomes[0].mutationsOfType(m2),individ
ual.genomes[1].mutationsOfType(m2))>=0];
                                Hetero = c(

        individual.genomes[0].mutationsOfType(m2)[match(individual.genomes[0].mutationsOfType(m2)
,individual.genomes[1].mutationsOfType(m2))<0],

        individual.genomes[1].mutationsOfType(m2)[match(individual.genomes[1].mutationsOfType(m2)
,individual.genomes[0].mutationsOfType(m2))<0]);

                                // checks continued
                                // print(Homo);
                                // print(Hetero);

                                // use phenotypic value to define individual tag
                                individual.tag = 
asInteger(sum(Homo.selectionCoeff)*10000)*2+asInteger(sum(Hetero.selectionCoeff*Hetero.mutationTy
```

```
pe.dominanceCoeff)*10000)*2; // multiply by 10000 since individual tag can only be integers
			}

			// if there is epistasis, define individual tags differently according to 
the epistasis scenario chosen
			else if (Epistasis == T)
			{
				individual.tag = 0; // first set individual tag to be 0
				for (EpistasisPair in 1:asInteger(NQTL/2)) // extract information 
on each epistatic pair for each individual
				{
					EpistasisMutation1 = 
individual.genomes.mutationsOfType(m2)[which(individual.genomes.mutationsOfType(m2).tag == 
2*EpistasisPair-1)];
					EpistasisMutation2 = 
individual.genomes.mutationsOfType(m2)[which(individual.genomes.mutationsOfType(m2).tag == 
2*EpistasisPair)];
					UniqueEpistasisMutation1 = unique(EpistasisMutation1);
					UniqueEpistasisMutation2 = unique(EpistasisMutation2);
					S1 = abs(UniqueEpistasisMutation1.selectionCoeff);
					S2 = abs(UniqueEpistasisMutation2.selectionCoeff);
					D1 = UniqueEpistasisMutation1.mutationType.dominanceCoeff;
					D2 = UniqueEpistasisMutation2.mutationType.dominanceCoeff;
					sign = (-1)^EpistasisPair; // give half of the pairs effect 
sizes the reverse of the epistatic scenario defined to make sure the mean trait value is ~0 in 
the first generation

					// for each pair, calculate its contribution to the 
phenotype and add it to the individual tag
					if (size(EpistasisMutation1) == 2)
					{
						//print(c("EpistasisMutation1Homo"));
						if (size(EpistasisMutation2) == 2)
	
	individual.tag=individual.tag+asInteger(EpiSce[8]/4*(2*S1 + 2*S2)*10000*sign);
						else if (size(EpistasisMutation2) == 1)
	
	individual.tag=individual.tag+asInteger(EpiSce[7]/3*(2*S1 + 2*D2*S2)*10000*sign);
						else
	
	individual.tag=individual.tag+asInteger(EpiSce[6]/2*(2*S1 + 0)*10000*sign);
					}
					else if (size(EpistasisMutation1) == 1)
					{
						//print(c("EpistasisMutation1Hetero"));
						if (size(EpistasisMutation2) == 2)
	
	individual.tag=individual.tag+asInteger(EpiSce[5]/3*(2*D1*S1 + 2*S2)*10000*sign);
						else if (size(EpistasisMutation2) == 1)
	
	individual.tag=individual.tag+asInteger(EpiSce[4]/2*(2*D1*S1 + 2*D2*S2)*10000*sign);
						else
	
	individual.tag=individual.tag+asInteger(EpiSce[3]/1*(2*D1*S1 + 0)*10000*sign);
					}
					else if (size(EpistasisMutation1) == 0)
					{
						if (size(EpistasisMutation2) == 2)
	
	individual.tag=individual.tag+asInteger(EpiSce[2]/2*(0 + 2*S2)*10000*sign);
						else if (size(EpistasisMutation2) == 1)
	
	individual.tag=individual.tag+asInteger(EpiSce[1]/1*(0 + 2*D2*S2)*10000*sign);
						else
							individual.tag=individual.tag;
					}
					else
					{
					print("ERROR");
					}
				}
```

```
                    }
        }

        // output the traits
        Traits = paste(sim.subpopulations.individuals.tag/10000, "\n"); //divide by 1000 to
return the original trait value
        if (Direction == F)
                writeFile(OutPath + "SimRep" + SimRepID + "/ExpRepPlus" + ExpRepID + "/Gen" +
GenID + "_Trait.txt", Traits, F);
        else
                writeFile(OutPath + "SimRep" + SimRepID + "/ExpRepMinus" + ExpRepID + "/Gen" +
GenID + "_Trait.txt", Traits, F);

        // output number of persisting QTLs each generation
        NSegregatingQTL =
paste(size(unique(sim.subpopulations.individuals.genomes.mutationsOfType(m2))));
        if (Direction == F)
                writeFile(OutPath + "SimRep" + SimRepID + "/ExpRepPlus" + ExpRepID + "/Gen" +
GenID + "_NSegregatingQTL.txt", NSegregatingQTL, F);
        else
                writeFile(OutPath + "SimRep" + SimRepID + "/ExpRepMinus" + ExpRepID + "/Gen" +
GenID + "_NSegregatingQTL.txt", NSegregatingQTL, F);

        // get the phenotypic order for all individuals in order to perform truncating selection;
change this if a different seelction regime is used
        Order = order(sim.subpopulations.individuals.tag, Direction);

        // redefine individual tags as their phenotypic ranks
        sim.subpopulations.individuals.tag[Order] = 0:(size(sim.subpopulations.individuals)-1);
}

10000: fitness(m2) {
        // the QTLs themselves are neutral; the m3 mutation type handles them
        return 1.0;
        }

10000: fitness(m3) {
        // keep only the 100 highest ranking individuals and remove all others
        if(individual.tag < SelectedSize)
                return 1.0;
        else
                return 0.0;
}

10000:10010 late() {
        GenID = sim.generation-9999;
        // last sanity check
        //print(GenID);
        //print(size(unique(sim.subpopulations.genomes.mutationsOfType(m1))));
        //print(size(unique(sim.subpopulations.genomes.mutationsOfType(m2))));
        //print(size(unique(sim.subpopulations.genomes.mutations)));
        AllIndividuals = sim.subpopulations.individuals;
        SampledIndividuals = sample(AllIndividuals, SampleSize);
        // output sampled population
        if (Direction == F)
                SampledIndividuals.genomes.output(OutPath + "SimRep" + SimRepID + "/ExpRepPlus" +
ExpRepID + "/Gen" + GenID + "_Sample.txt");
        else
                SampledIndividuals.genomes.output(OutPath + "SimRep" + SimRepID + "/ExpRepMinus" +
ExpRepID + "/Gen" + GenID + "_Sample.txt");

        // output full population
        if (Direction == F)
                AllIndividuals.genomes.output(OutPath + "SimRep" + SimRepID + "/ExpRepPlus" +
ExpRepID + "/Gen" + GenID + "_Full.txt");
        else
                AllIndividuals.genomes.output(OutPath + "SimRep" + SimRepID + "/ExpRepMinus" +
ExpRepID + "/Gen" + GenID + "_Full.txt");
}
```

### Figures

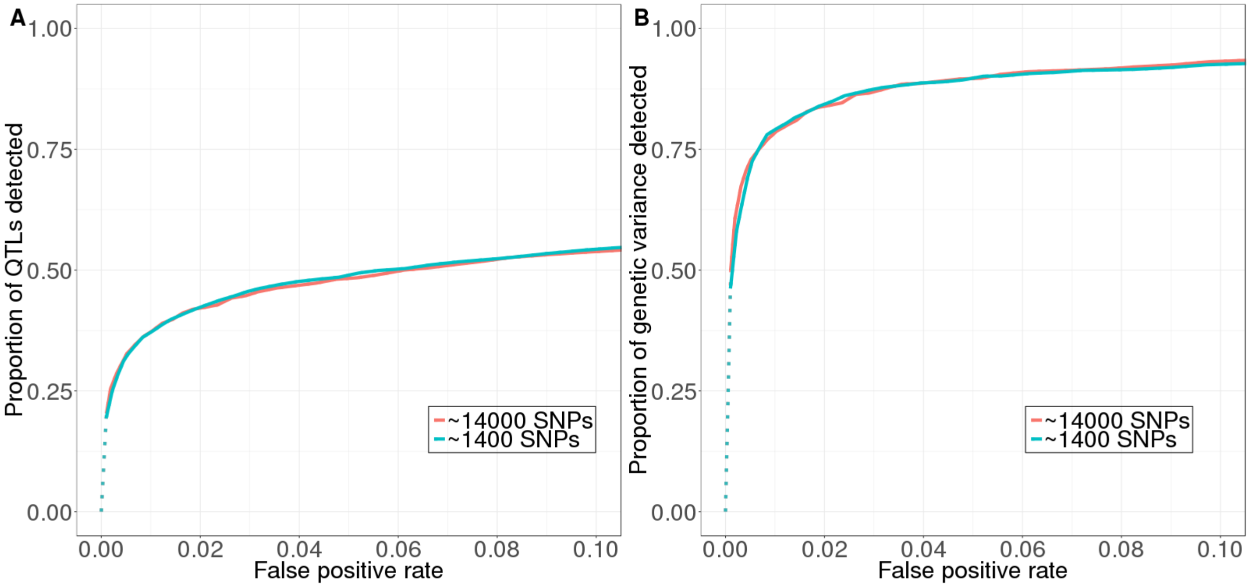

**Figure S1.** Number of SNPs in the simulated population has little effect on the power and false positive rate in QTL detection. (A) Power measured by the proportion of QTL detected. (B) Power measured by the proportion of genetic variance in the first generation explained by the detected QTL. Our standard model has ~14,000 SNPs, and the model used to test computational methods has ~1,400 SNPs. *D*-value is used here to detect QTLs.

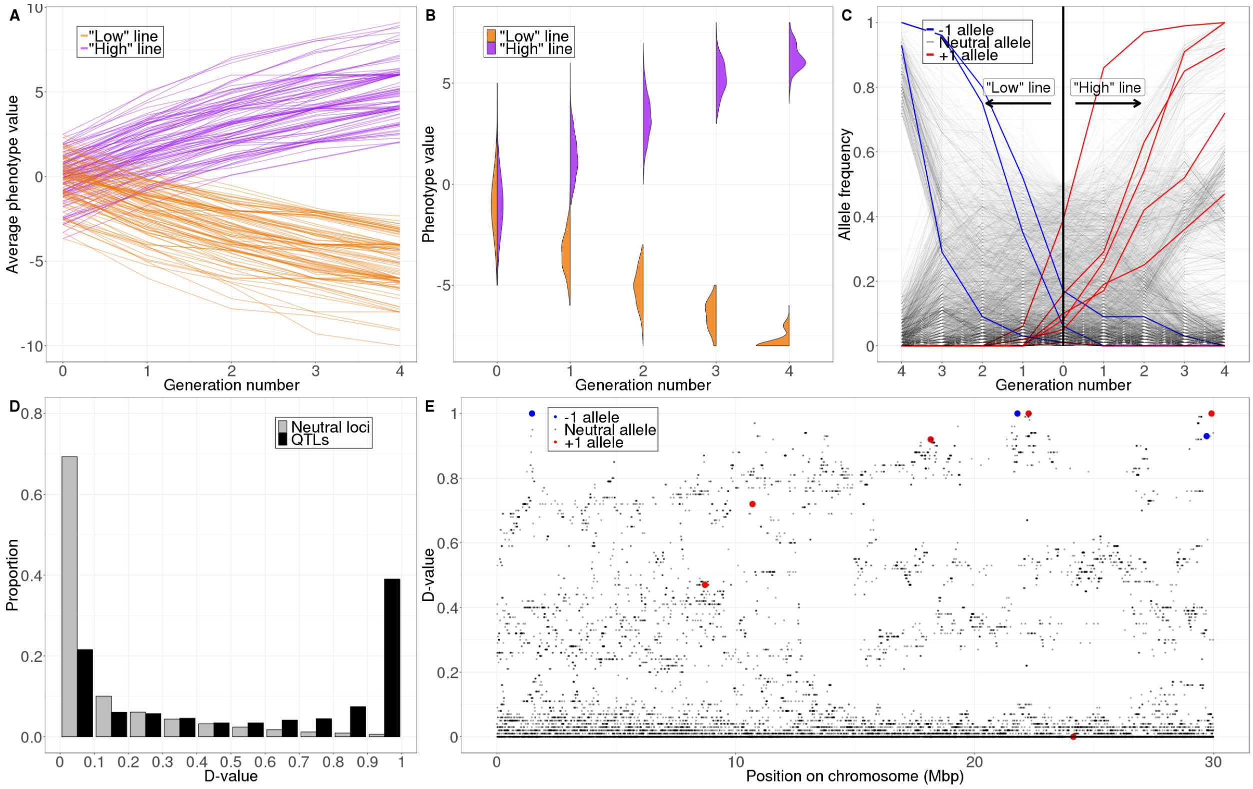

**Figure S2.** Simulation result under our standard model with 10 QTLs. (A) Change in the average phenotype values in all 100 simulation replicates. Each line represents a selection line in one simulation replicate. (B) Change in the distribution of trait values in the population over one single simulation run. (C) Change in sampled minor allele frequencies at neutral loci and QTLs in one single simulation run. The left half of the figure shows the low line and the right half shows the high line. (D) Distribution of *D*-values per locus across all 100 simulation replicates grouped by neutral loci (grey bars) vs QTLs (black bars). (E) *D*-values of neutral loci and QTLs along the simulated chromosome in one single simulation run.

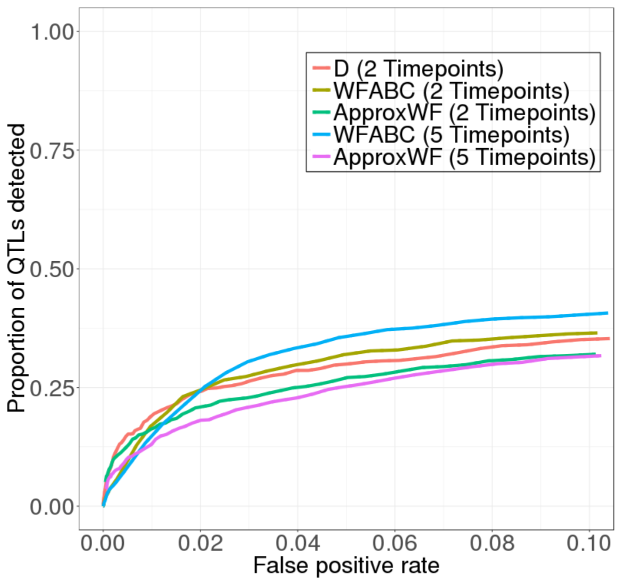

**Figure S3** Performance of different detection methods when only one selection line is created. In this scenario, WFABC gains an advantage over D at higher false positive rates. A slightly modified version of the standard model with reduced level of nucleotide diversity and 10 QTLs is used in this comparison (same model as shown in the blue line in Figure S1. Power is measured by the proportion of QTLs detected.

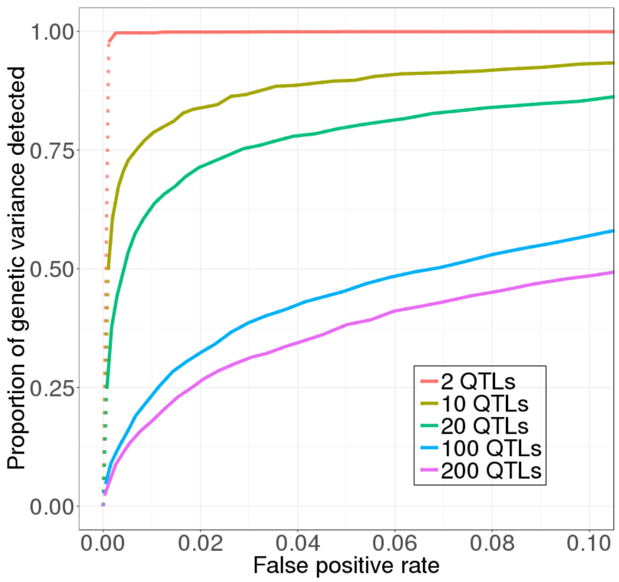

**Figure S4.** Detection power is lower when more QTL underlie the trait under selection. The standard QTL model is used in this comparison. Power measured by the proportion of genetic variance in the first generation explained by the detected QTL.

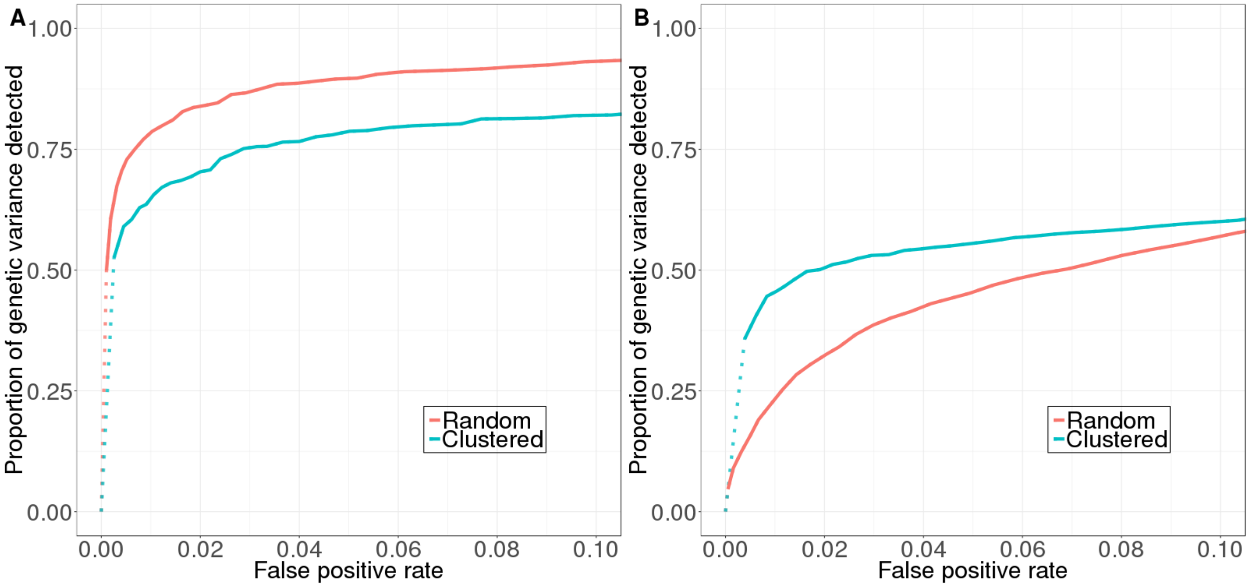

**Figure S5.** The clustering of QTL has different effects on detection power depending on the number of QTL underlying the trait under selection. (A) With 10 QTL, clustering reduces detection power. (B) With 100 QTL, clustering increases detection power at lower false positive rate. Power measured by the proportion of genetic variance in the first generation explained by the detected QTL.

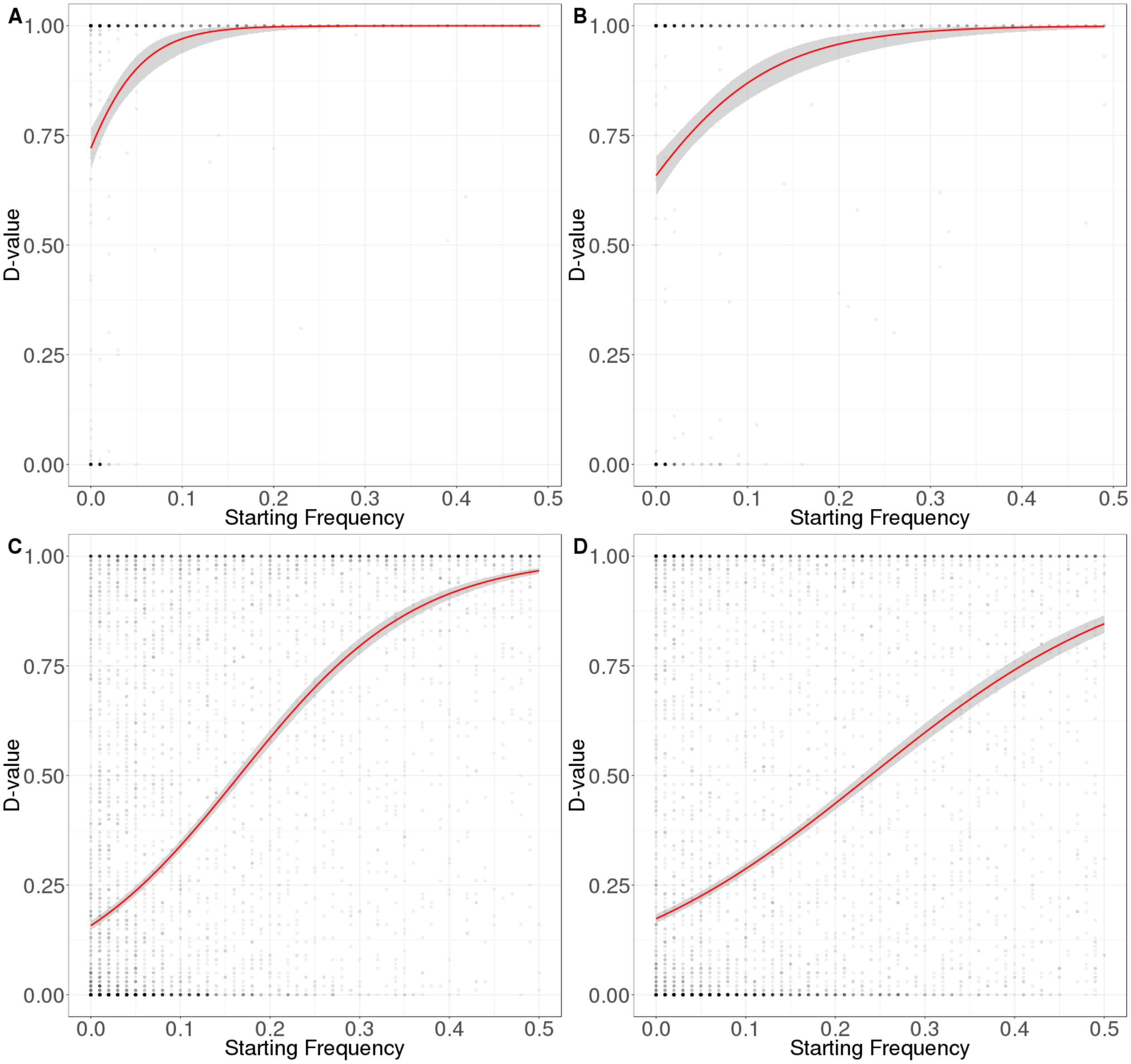

**Figure S6.** (A) 10 QTLs, equal effect sizes (B) 10 QTLs, exponential effect sizes (C) 100 QTLs, equal effect sizes (D) 100 QTLs, exponential effect sizes.

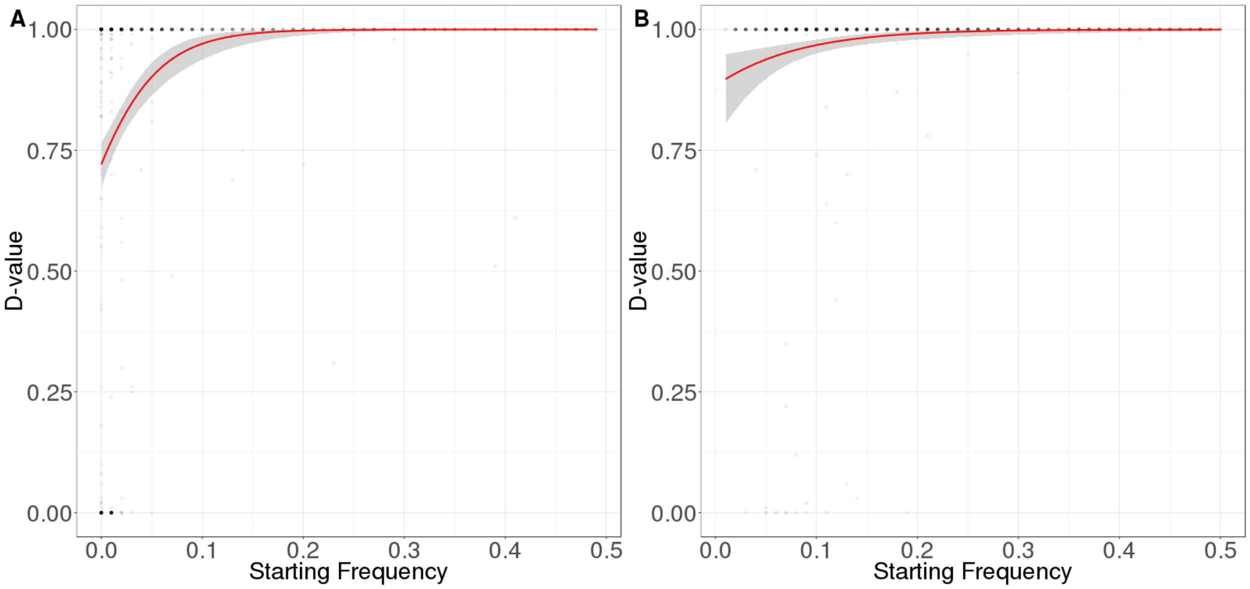

**Figure S7** Relationship between *D* and starting frequencies of minor alleles at QTL across all 100 simulation replicates. A logistic regression model is fit to the data, shown with the red line. (A) Random starting frequency. (B) High starting frequency (>5%). When QTL start at a random frequency, the QTL that can be easily detected (i.e. those with high D-values) are more likely to start at higher frequencies. Therefore, they contribute more to the variance in the first generation. A model with 10 QTL is shown in this figure.

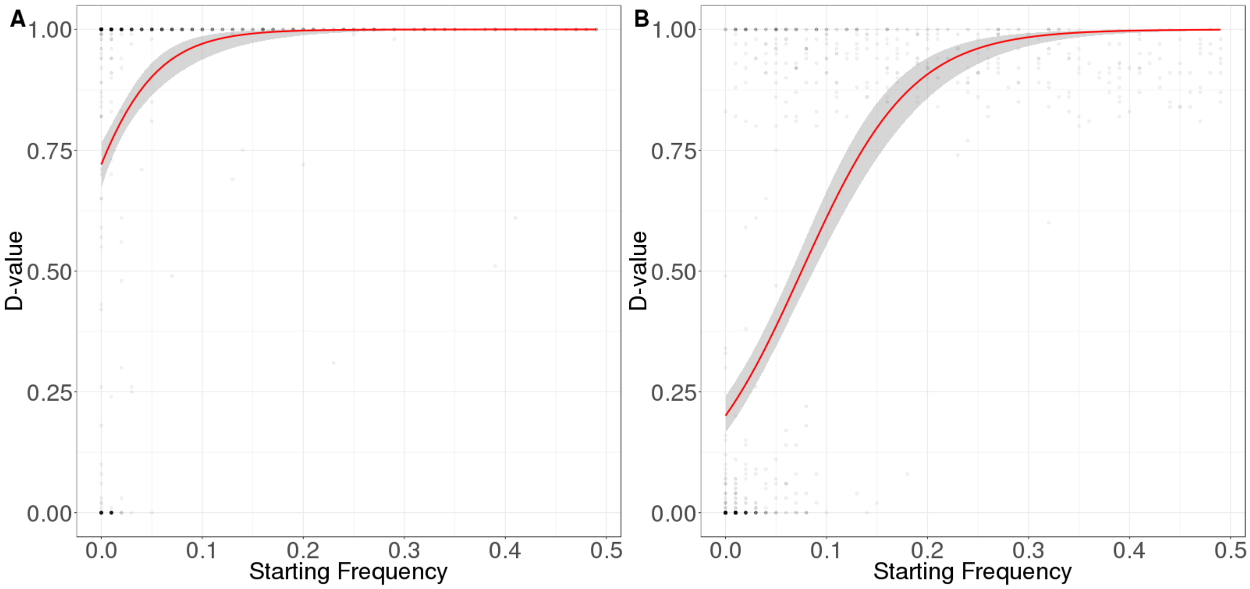

**Figure S8** Relationship between *D* and starting frequencies of minor alleles at QTL across all 100 simulation replicates. A logistic regression model is fit to the data, shown with the red line. (A) Mutant is codominant. (B) Mutant is recessive. When mutant is recessive, the QTL that can be easily detected (i.e. those with high D-values) are more likely to start at higher frequencies. Therefore, they contribute more to the variance in the first generation.

### Tables

**Table S1** Pairwise epistasis scenarios. Lowercase letters (a and b) represent the wild type alleles, and uppercase letters (A and B) represent the mutant alleles. Numerical entries are the contributions to the phenotype under selection by given genotype combinations. The signs of these contributions are randomly assigned. These contributions are additive among all epistatic pairs.

Synergistic, weak

| | bb | Bb | BB |
|---|---|---|---|
| aa | 0 | 1 | 2 |
| aA | 1 | 2.5 | 4 |
| AA | 2 | 4 | 6 |

Synergistic, strong

| | bb | Bb | BB |
|---|---|---|---|
| aa | 0 | 1 | 2 |
| aA | 1 | 3 | 5 |
| AA | 2 | 5 | 8 |

Antagonistic, weak

| | bb | Bb | BB |
|---|---|---|---|
| aa | 0 | 1 | 2 |
| aA | 1 | 1.5 | 2 |
| AA | 2 | 2 | 2 |

Antagonistic, strong

| | bb | Bb | BB |
|---|---|---|---|
| aa | 0 | 1 | 2 |
| aA | 1 | 1 | 1 |
| AA | 2 | 1 | 0 |

Sign, weak

| | bb | Bb | BB |
|---|---|---|---|
| aa | 0 | 1 | 2 |
| aA | -1 | 1 | 3 |
| AA | -2 | 1 | 4 |

Sign, strong

| | bb | Bb | BB |
|---|---|---|---|
| aa | 0 | 1 | 2 |
| aA | -1 | 1.5 | 4 |
| AA | -2 | 2 | 6 |

Reciprocal sign, weak

| | bb | Bb | BB |
|---|---|---|---|
| aa | 0 | -1 | -2 |
| aA | -1 | 0 | 1 |
| AA | -2 | 1 | 4 |

Reciprocal sign, strong

| | bb | Bb | BB |
|---|---|---|---|
| aa | 0 | -1 | -2 |
| aA | -1 | 0.5 | 2 |
| AA | -2 | 2 | 6 |
